# Regulating IL-2 immune signaling function via a core allosteric structural network

**DOI:** 10.1101/2024.10.07.617024

**Authors:** Claire H. Woodward, Shahlo O. Solieva, Daniel Hwang, Viviane S. De Paula, Charina S. Fabilane, Michael C. Young, Tony Trent, Ella C. Teeley, Ananya Majumdar, Jamie B. Spangler, Gregory R. Bowman, Nikolaos G. Sgourakis

## Abstract

Human interleukin-2 (IL-2) is a crucial cytokine for T cell regulation, with therapeutic potential in cancer and autoimmune diseases. However, IL-2’s pleiotropic effects across different immune cell types often lead to toxicity and limited efficacy. Previous efforts to enhance IL-2’s therapeutic profile have focused on modifying its receptor binding sites. Yet, the underlying dynamics and intramolecular networks contributing to IL-2 receptor recognition remain unexplored. This study presents a detailed characterization of IL-2 dynamics compared to two engineered IL-2 mutants, “superkines” S15 and S1, which exhibit biased signaling towards effector T cells. Using NMR spectroscopy and molecular dynamics simulations, we demonstrate significant variations in core dynamic pathways and conformational exchange rates across these three IL-2 variants. We identify distinct allosteric networks and excited state conformations in the superkines, despite their structural similarity to wild-type IL-2. Furthermore, we rationally design a mutation (L56A) in the S1 superkine’s core network, which partially reverts its dynamics, receptor binding affinity, and T cell signaling behavior towards that of wild-type IL-2. Our results reveal that IL-2 superkine core dynamics play a critical role in their enhanced receptor binding and function, suggesting that modulating IL-2 dynamics and core allostery represents an untapped approach for designing immunotherapies with improved immune cell selectivity profiles.

**Highlights:**

- NMR and molecular dynamics simulations revealed distinct conformational dynamics and allosteric networks in computationally re-designed IL-2 superkines compared to wild-type IL-2, despite their similar crystal structures.
- The superkines S1 and S15 exhibit altered sampling of excited state conformations at an intermediate timescale, with slower conformational exchange rates compared to wild-type IL-2.
- A rationally designed mutation (L56A) in the S1 superkine’s core allosteric network partially reverted its dynamics, receptor binding affinity, and T cell signaling behavior towards that of wild-type IL-2.
- Our study demonstrates that IL-2 core dynamics play a critical role in receptor binding and signaling function, providing a foundation for engineering more selective IL-2-based immunotherapies.

**Graphical Abstract:** 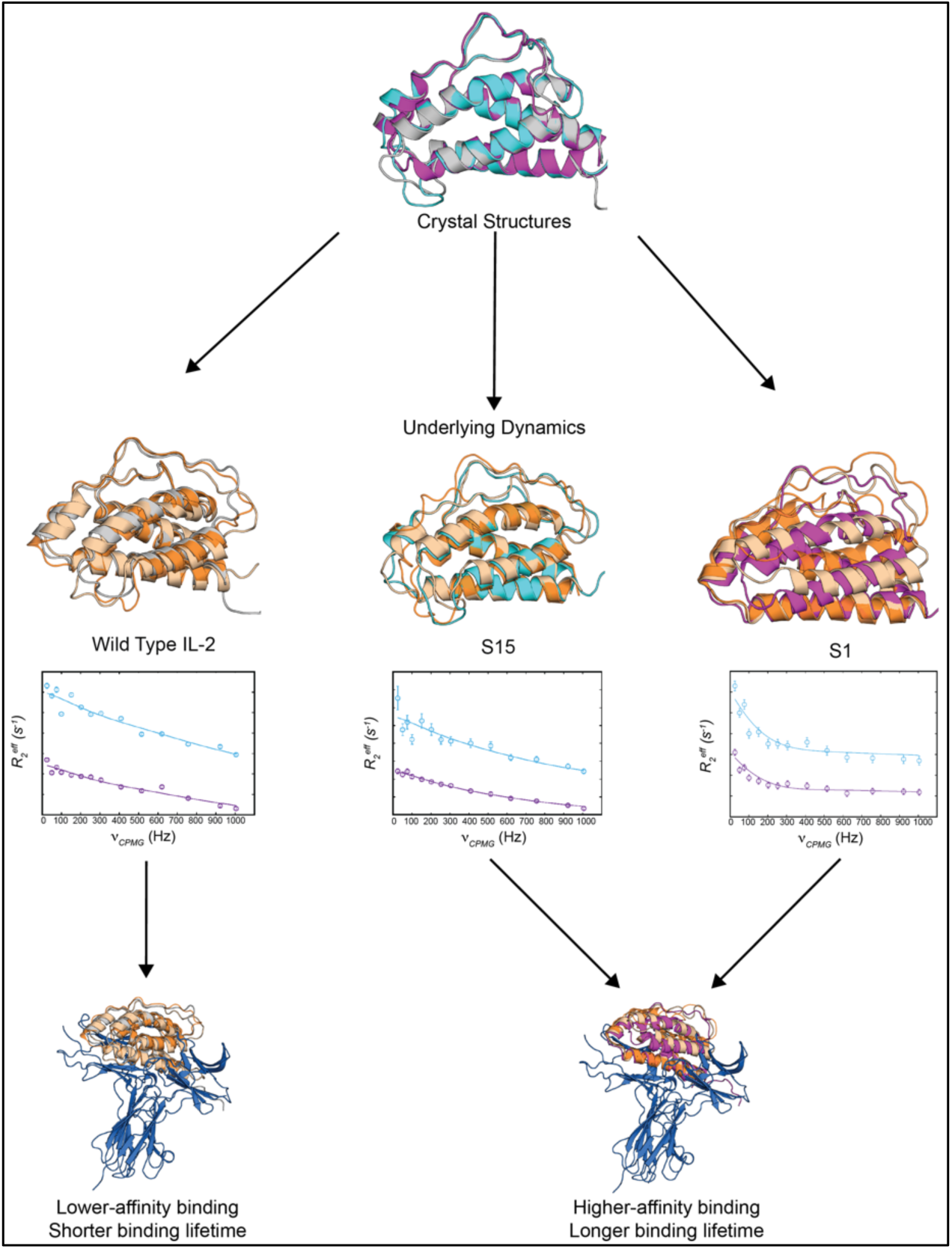

## Introduction

Interleukin-2 (IL-2), is a cytokine in the common gamma chain family that promotes essential functions of immune cells including expansion, differentiation, and maintenance^1,2^. In particular, its role in T lymphocyte regulation and proliferation has been characterized extensively^3,4^. IL-2 signals on T cells via two receptor (IL-2R) forms: an intermediate-affinity heterodimeric receptor, comprised of IL-2Rβ (CD122) and γ_c_ (CD132) subunits, or a high-affinity heterotrimeric receptor complex comprised of IL-2Rα (CD25), IL-2Rβ (CD122), and γ_c_ (CD132) subunits^5–7^. Thus, differences in IL-2Rα expression across T cell subsets result in distinct IL-2 signaling potencies across cell types^6^. Specifically, Foxp3^+^ T regulatory (T_reg_) cells express high levels of IL-2Rα whereas naïve effector T (T_eff_) cells and natural killer (NK) cells express low levels of IL-2Rα^6^. Because IL-2 plays a central role in essential T cell-mediated immunity, it is a desirable target for therapeutics^8,9^. Recombinant human IL-2-based therapy is administered in high doses to treat cancers such as malignant melanoma and renal cell carcinoma and aims to stimulate the intermediate-affinity dimeric IL-2R found on tumor-infiltrating CD4^+^ and CD8^+^ T cells and natural killer cells^10^. Alternatively, Foxp3^+^ T regulatory (T_reg_) cells are highly sensitive to IL-2 due to their distinctively high levels of IL-2Rα expression^2^; therefore, low-dose IL-2 therapies have also undergone clinical testing for the treatment of multiple autoimmune diseases^11–14^. In either case, clinical use is limited by IL-2’s ability to simultaneously induce multiple, conflicting types of immune responses through activation of several cell types (pleiotropy), leading to low efficacy of these treatments and systemic toxicity in patients^15–17^. In extreme situations, high-doses of IL-2 treatment can target IL-2Rα^+^ endothelial cells, leading to the release of vasoactive mediators which cause increased capillary permeability, leading to vascular leak syndrome^18–20^. Previous efforts to improve clinical translation focused on engineering IL-2 mutants by manipulating the cytokine’s direct interaction surfaces with IL-2R subunits^21–32^. However, particularly in the development of autoimmune disease treatments, existing T_reg_-biasing strategies rely heavily on differentials in IL-2Rα expression across T cell subsets (which vary between patients), leading to significant limitations in using binding interface mutations to target T_reg_ cells for specific activation. This is best explained by two key features of the system: *i)* IL-2Rα is not required for IL-2 binding or signaling, so increasing the binding affinity for IL-2Rα alone has minimal T_reg_-selective biasing impact, and *ii)* both IL-2Rβ and γ_c_ are required for signal transduction, so decreasing the affinity of IL-2 for the IL-2Rβγ complex (in order to increase reliance on IL-2Rα) effectively antagonizes the IL-2R signaling function^24,29–36^.

An alternative approach would be to design IL-2 functional mimetics that favor or disfavor interactions with specific receptor subunits without directly altering their native interaction surfaces on IL-2. Previous structural studies revealed that there are no direct interactions between IL-2Rα with the other receptor subunits^7,37^. However, comparison of multiple X-ray structures suggests that IL-2Rα binding to IL-2 induces a conformation that is more favorable for interactions with IL-2Rβ^7,38,39^. These findings support a model where allosteric structural changes allow for positive cooperativity between the IL-2Rα and IL-2Rβ binding interfaces on opposite sides of the IL-2 structure. This model is further supported by an engineered superkine (denoted H9), a functional IL-2 mutant isolated through directed evolution, which has enhanced binding to IL-2Rβ and more potent signaling through the dimeric receptor (IL-2Rβγ_c_) expressed on resting T_eff_ cells^21^. The six superkine mutations are mostly removed from the receptor binding interface, and have been suggested to lock the C-helix into a conformation resembling the IL-2Rα-bound structure of IL-2 thereby bypassing reliance on IL-2Rα for high-affinity interactions with IL-2Rβ^21^. Furthermore, recent protein engineering advancements led to the development of computationally redesigned IL-2 superkines with equally enhanced T_eff_ signaling outcomes^22^. These superkine designs show significantly enhanced protein stability through improved side chain packing, loop remodeling and other structural remedies of the native IL-2 structure, leading to an approximately 5-15-fold improved affinity for IL-2Rβ. Interestingly, the redesigned superkine mutations are also generally removed from the IL-2Rβ binding site on IL-2, and crystal structures demonstrate high structural similarity to wild type IL-2^22^. While these findings suggest a more rational approach for enhancing IL-2’s function through structure-guided design, the specific network of core interactions and structural transitions that mediate such functional outcomes remains incompletely characterized.

To further understand IL-2 cytokine/receptor interactions and how they may be manipulated to bias immune behavior, we studied its dynamic conformational transitions along with their functional significance in terms of interactions with IL-2R receptor subunits and signaling function using an integrative approach bridging NMR spectroscopy, surface plasmon resonance (SPR), molecular dynamics (MD) simulations, and in vitro T cell activation assays. Focusing on two engineered IL-2 variants (superkines S1 and S15), we identified key structural elements and elucidated their dynamic conformational transitions that regulate their function. Notably, altering a linchpin residue of the S1 superkine’s core allosteric network reduced its receptor binding and T_eff_ cell signaling activity. Our findings highlight the importance of IL-2’s structural flexibility in receptor interactions, providing a foundation for engineering more effective IL-2-based immunotherapies that could target specific immune cell subsets with greater precision.

## Results

### NMR chemical shift mapping reveals changes in core packing environment for engineered IL-2 variants

NMR can provide a high-resolution view of internal and global protein motions and glean insights into subtle structural differences between different IL-2 variants in a solution environment^40^. Thus, we prepared isotopically labelled protein samples and carried out backbone and methyl resonance assignments for the S1 and S15 superkines, to complement our previous assignments of WT human IL-2^41^. Robust assignments at high levels (89%/95% and 92%/97% for the backbone/methyl atoms of S15 and S1, respectively), were accomplished using a combination of triple resonance experiments and amide-methyl and methyl-methyl NOESY experiments, further assisted by our previously published human IL-2 assignments (Fig. S1 and S2)^41^. Our backbone assignments are in agreement with the existing structural data, confirming that the overall secondary structure and backbone conformation is similar across the wild type and superkine proteins, based on structural predictions from chemical shifts (Fig. S2A-B)^42,43^. Furthermore, methyl-methyl NOE connectivities recorded using selectively ^13^C-labeled methyl side chains of isoleucine, leucine, and valine residues confirm that the average solution conformation of each superkine closely aligns with its respective crystal structure (Figure S2C-D).

To identify distinguishing features in different IL-2 variants, we directly compared the NMR spectra of each superkine relative to wild type IL-2 (Fig. 1). To represent 2D ‘fingerprints’ of the amide backbone and methyl side chain residues, we overlaid transverse relaxation-optimized spectroscopy (TROSY) and ^13^C-^1^H band-selective optimized flip angle short transient (SOFAST) heteronuclear multiple quantum coherence (HMQC) spectra. This enables mapping of the effects of the mutations on the backbone and hydrophobic core of each IL-2 variant (Fig. 1A-B). Given that there are 7 and 17 amino acid substitutions between S15 and S1 relative to wild type IL-2, respectively (Fig. 1C), we expected that significant differences in the local chemical environment would be primarily localized near the mutation sites. To quantify this, we calculated the chemical shift perturbations (CSPs) observed in the 2D spectra to monitor structural differences in the backbone amide residues and the methyl sidechains composing the hydrophobic core (Fig. 1D-E, S2E-F). We observed that the largest CSPs were at or near the mutation sites, as expected. However, when we mapped the methyl CSPs onto the S1 and S15 crystal structures, we detected significant long-range CSPs in the hydrophobic core of each protein. Thus, although the three proteins have nearly identical crystal structures (Fig. S2D) and similar backbone conformations in solution, methyl NMR enables detection of minor changes in the average solution environments of core sidechains.

**Figure 1:**
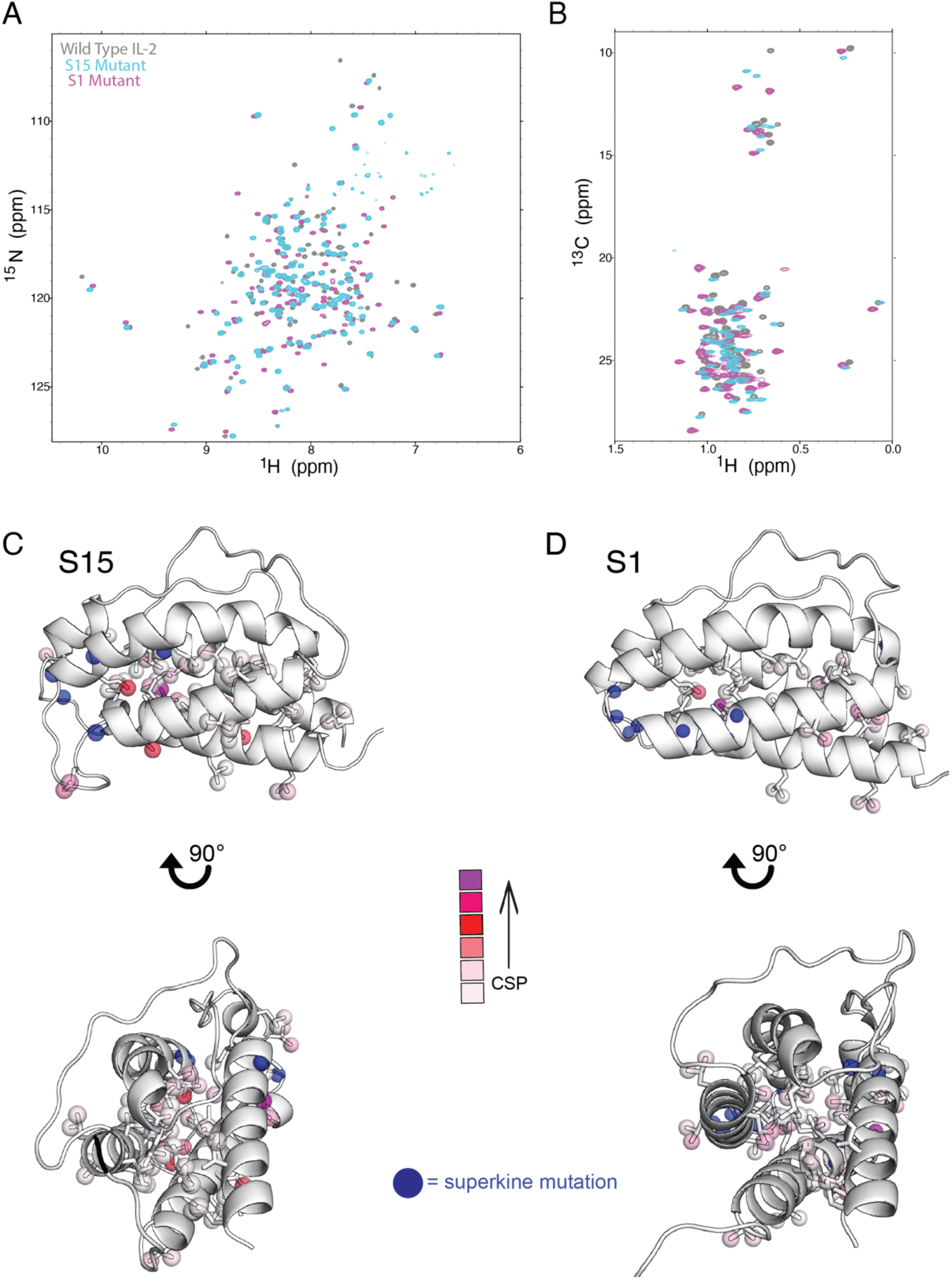
NMR spectra highlight local changes between wild type IL-2 and superkines in a solution environment. A) ^15^N-^1^H TROSY HSQC and B) ^13^C-^1^H methyl SOFAST HMQC spectra overlays of wild type IL-2 (gray), S15 superkine (cyan), and S1 superkine (magenta). All spectra were collected at 800 MHz ^1^H magnetic field and 25°C. Methyl chemical shift perturbations (CSPs) plotted as spheres in a color gradient on the C) S15 and D) S1 crystal structures (PDB IDs: 7raa and 7ra9). Superkine mutation sites on each protein are plotted as blue spheres for reference.

### Fast time scale dynamics show differences in backbone rigidity between IL-2 and the superkines

To detect differences in fast motions of the IL-2 backbone between wild type IL-2, S15, and S1, we assessed each protein using model-free analysis of transverse (R_2_) and longitudinal (R_1_) ^15^N relaxation rate constants as well as heteronuclear NOE values at two magnetic fields (Fig. S3)^44,45^. The experimental data were then analyzed using the model-free formalism via the program Relax^46–49^ to extract order parameters (S^2^), which describe the degree of restriction of the motions of amide bond vectors in the molecular frame (Fig. S4 & S5). S^2^ values ranging from 0 to 1 indicate a bond vector that is completely unrestricted or restricted, respectively, on a molecular frame defined by the rotational diffusion of the molecule. We observed that wild type IL-2 and S15 had similar diffusion tensors and rotational correlation times (defined as the average time required for a 1 radian rotation), whereas S1 demonstrated distinct diffusion properties with a slightly faster rotational correlation time and slightly different diffusion model. As relaxation rates are impacted by dimerization, we confirmed that this could not be explained by concentration-dependent dimerization by detecting R_2_ relaxation rates with a diluted sample (Fig. S4). One explanation is that shortening and stabilizing the “floppy” BC-loop results in reduced “drag” in solution, yielding an altered diffusion model and slightly faster rotational correlation time^22^. Close inspection of the order parameters for all three proteins reveals that the alpha-helical regions of the structure are relatively rigid (order parameter >0.8). Flexible regions generally correspond to connecting loops; most notable is the CD-loop, which has extremely low order parameters (<0.5) at the N-terminal region of the loop (resid. ∼99-104) and moderately low (0.6-0.8) on the C-terminal half (resid. ∼105-112), likely controlled by the disulfide bridge connecting this loop to the B-helix (resid. C105 & C58) (Fig. S4). These general features apply to all three proteins, indicating that the overall backbone characteristics agree as expected for proteins with very similar secondary structures.

Comparison of the order parameters obtained for wild type IL-2 relative to the two superkines reveals several key differences in picosecond-nanosecond backbone motions (Fig. 2). Each superkine displays distinct regions where fast timescale motions are altered; positive and negative changes are spread throughout the S15 and S1 structures, indicating that impacts of the mutations are quite complex and far-reaching. In S15, we observe that the ends of alpha helices are more rigid relative to wild type IL-2, and solvent-exposed loop residues are generally more flexible (Fig. 2A & 2C). In S1, we observe a different trend: several solvent-exposed residues in the A- and C-helices are more rigid in S1 compared to wild type IL-2 (Fig. 2B & 2D). This aligns with the design principles of S1, given that it was designed to have a stabilized C-helix due to several mutations in the BC-loop^22^. Notably, residues in the A- and C-helices form the interface with IL-2Rβ^7,37^, so this rigidification of S1 may explain the enhanced receptor binding characteristic of superkines (Fig. 2E). S15, which does not contain the remodeled BC-loop design, shows minimal detectable changes along these helices. Another important region is the mini (A’)-helix, which is more rigid on the solvent-exposed side of the central turn in S1, whereas the ends of this helix are more flexible compared to wild type IL-2 (Fig. 2B). Changes in the dynamics of this region are not surprising given the two nearby mutations (resid. 32 & 39), but these effects could conceivably contribute to the previously described reduction in IL-2Rα binding, in combination with the disruptive L72A mutation^22^ (Fig. 2E). We were unable to confidently assign the mini-helix amides in S15 for comparison, which in itself may point toward the presence of exchange broadening effects in this region, also an indicator of altered backbone dynamics. Taken together, it is clear that S15 and S1 each demonstrate widespread, distinctive backbone dynamics on the fast timescale as a result of the superkine mutations.

**Figure 2:**
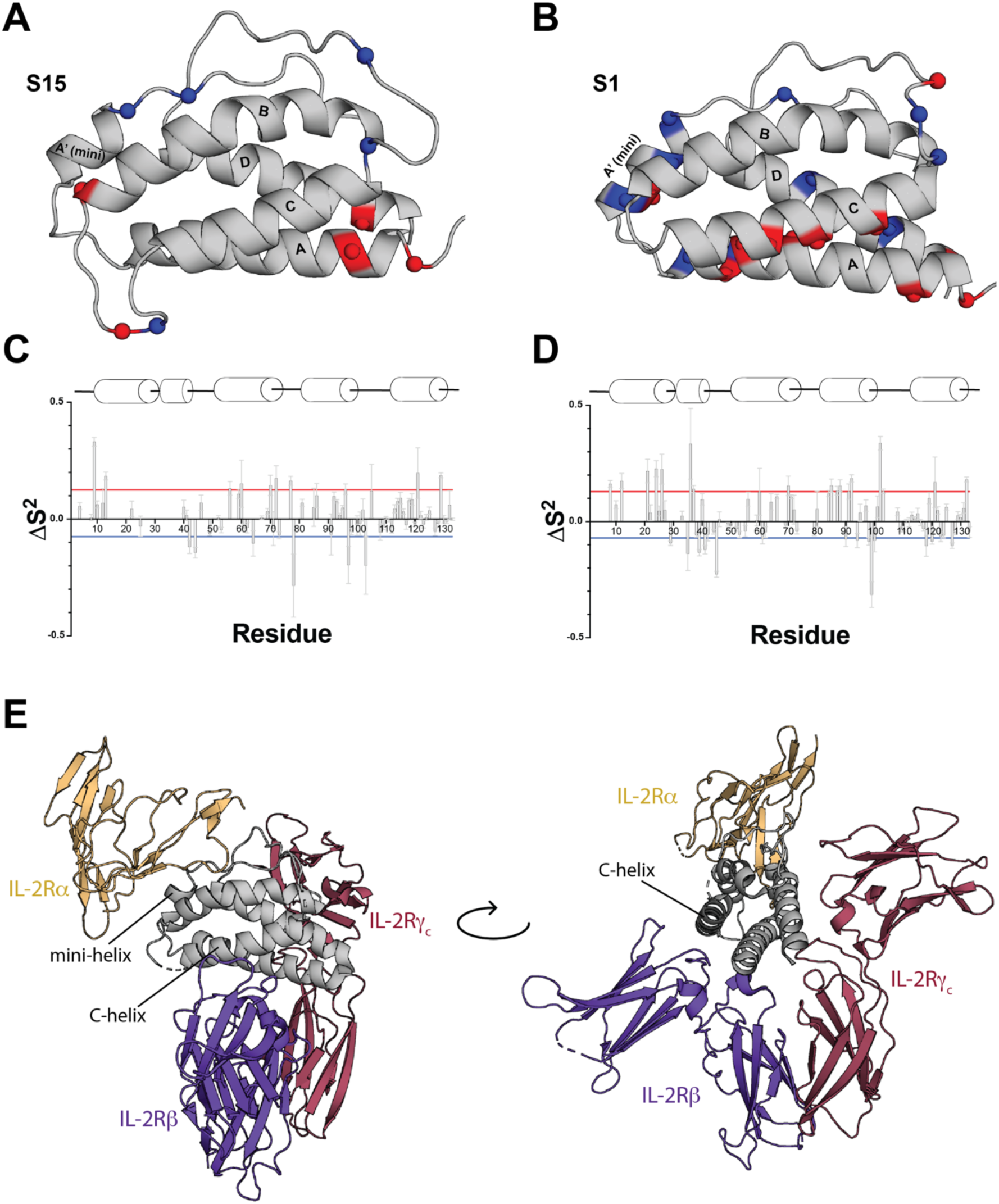
Superkines show altered backbone rigidity on the picosecond-nanosecond timescale relative to wild type IL-2. Significant differences in order parameters (ΔS^2^) relative to wild type IL-2 are plotted on the A) S15 (PDB ID: 7raa) and B) S1 (PDB ID: 7ra9) structures as blue (negative ΔS^2^ = increased flexibility) or red (positive ΔS^2^ = increased rigidity) spheres. Raw ΔS^2^ values are shown as a bar graph for C) S15 and D) S1, with secondary structure shown above for reference. Significance is defined as ΔS^2^ at least 10% above or below the mean, shown as red and blue boundaries. S1 residue numbering is based on alignment to the wild type sequence. E) Crystal structure of IL-2 in complex with the heterotrimeric IL-2R, included for reference (PDB ID: 2erj).

### The superkines exhibit altered sampling of excited state conformations at an intermediate time scale

Given our observation of subtle differences in the hydrophobic core environment between IL-2 and its superkine variants (Fig. 1), we hypothesized that subtle changes in conformations adopted by methyl-bearing sidechains may be important for receptor binding. To further characterize differences in dynamics occurring at an intermediate (microseconds-to-milliseconds timescale), we used methyl-selective Carr-Purcell-Meiboom-Gill (CPMG) relaxation dispersion experiments^50^. When conformational exchange occurs on the microsecond to millisecond timescale by NMR, a peak broadening effect is observed which is quantifiable by monitoring the so-called ‘effective transverse relaxation rate’ (R_2_^eff^). In the CPMG relaxation dispersion experiment, a series of ‘refocusing’ pulses are used to quench the peak broadening effects, which are inversely proportional to R_2_^eff^. Therefore, as the frequency of CPMG refocusing pulses increases, we expect to observe a decrease in the R_2_^eff^ values if conformational exchange is present (Fig. 3A). We observed multiple residues undergoing conformational exchange, suggesting the sampling of minor states. We then fitted our dispersion data using a global two-site model for conformational exchange, to extract the rate of exchange (k_ex_), population of the minor state (p_B_), and the chemical shift differences between the major and minor states (|ω|)^51^. We fitted a global exchange rate, k_ex_ of 4088 ± 190 s^-1^ for wild type IL-2, which is at the fast detection limit of the CPMG experiment. In comparison, both S15 and S1 display significantly slower rates of conformational exchange, at 2878 ± 44 s^-1^ and 651 ± 16 s^-1^, respectively. While our fits of global p_B_ values are insufficient to provide quantitative population differences across the three proteins, due to degeneracy in the χ^2^ landscape, a systematic transition towards lower global k_ex_ fit values can be observed for S1 relative to S15 and WT IL-2 (Fig. S6). These results indicate that, compared to wild type, the superkines experience slower transitions between the major and minor states.

**Figure 3:**
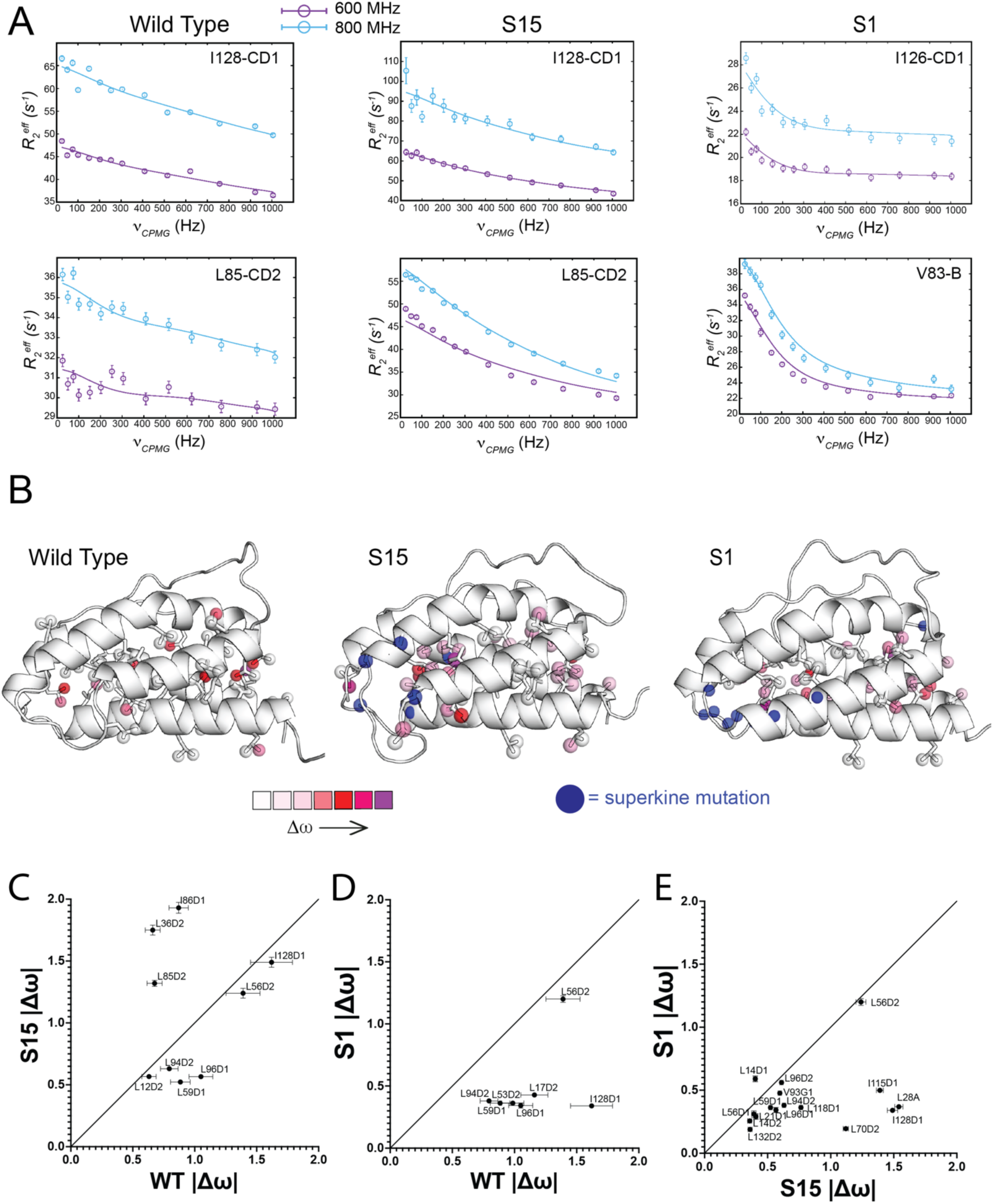
Superkines show perturbed intermediate-timescale dynamics relative to wild type IL-2. A) Representative dispersion curves extracted for wild type (left), S15 (middle) and S1 (right) proteins from methyl-selective, single-quantum CPMG relaxation dispersion experiments collected at 600 (purple) and 800 (cyan) MHz ^1^H magnetic fields at 5°C. Shown are dispersion curves for Ile128-CD1 and Leu85-CD2 (note that S1 has a two amino acid deletion in the BC-loop, so numbering is offset above residue 80 in the sequence). Experimental data are shown as circles in all panels, with errors estimated from the S/N in the raw experimental data. The best-fit lines are shown for a global analysis using a two-site conformational exchange model of all methyls with relaxation dispersion profiles showing Rex > 3, where Rex is defined as R2^eff^ (25 Hz) – R2^eff^ (1005 Hz). B) Methyl probes undergoing conformational exchange by CPMG are colored spheres on the model structures of wild type IL-2 (left, PDB ID: 1m47), S15 (middle, PDB ID: 7raa), and S1 (right, PDB ID: 7ra9) and colored as a gradient according to the magnitude of |Δω| values from the global two-site exchange fit. For reference, the mutation sites on S15 and S1 are shown as blue spheres. X vs. y correlation plots of |Δω| values from the global two-site exchange fit plotted as C) S15 vs. wild type IL-2, D) S1 vs. wild type IL-2, and E) S1 vs. S15, with names of methyl probes shown next to each point. S1 residue numbering is based on alignment to the wild type sequence.

Conformational exchange is observed for several methyl resonances throughout the wild type, S15, and S1 structures, although the specific profiles of the three proteins are different (Fig. 3, S6-10, Table S1). Significant differences in the global dynamic landscapes of these proteins are supported by 1) specific residues involved in each process differ across the three proteins, and 2) quantifiable per-residue structural ranges of motion between major and minor state (|Δω|) (Fig. 3B, S6). Specifically, we note that wild type IL-2 has both fewer residues undergoing exchange and a slightly lower R_ex_ (defined here as R_ex_ = R_2_^eff^(25 Hz) – R_2_^eff^(1005 Hz)) baseline overall compared to both superkines. Although S1 also has a slightly higher R_ex_ baseline compared to wild type IL-2, this is potentially a side effect of the differences in exchange rates (k_ex_) of the processes observed in each protein: the wild type IL-2 exchange rate is at the fast limit of the detection window by CPMG relaxation dispersion, and the S1 exchange rate is much slower and thereby easier to detect by CPMG, possibly causing the R_ex_ rates for S1 to be artificially slightly higher (Fig. S10). This is unlikely to be the case for S15, which retains a faster k_ex_ compared to the S1 superkine. Interestingly, the S15 superkine has two clusters of R_ex_ values – a cohort of lower R_ex_ methyls that have similar magnitude to wild type IL-2 and S1, and a cohort of high R_ex_ methyls with unique behavior (Fig. S10).

In each CPMG dataset, we included the methyl residues undergoing significant conformational exchange (defined by R_ex_ > 3 s^-1^) and fit the dispersion profiles using a global two-site exchange model, omitting residues with poor signal to noise or spectral overlap (Table S1, Fig. S6). Fitted |Δω| values are compared across the three structures to determine whether the conformation of the minor state is similar or different across different proteins. Here, we note that there are a handful of residues that are exclusively exchanging in each protein, preventing a comprehensive structural comparison from |Δω| values alone. Among the residues that can be compared in correlation plots, we observe that there is a cluster of residues that tend to correlate quite well across proteins, including residues in the A-helix, B-helix, and C-terminal end of the C-helix (Fig. 3C-E). When comparing the S15 superkine to wild type IL-2, we observe that residues in the mini (A’)-helix and N-terminal end of the C-helix have distinct behavior (Fig. 3C). This is an intriguing result given that these regions have been shown to undergo conformational changes upon binding to IL-2Rα or IL-2Rβ^7,21,22,37,39^. Comparing the S1 superkine to wild type IL-2, we observe that the overall magnitudes of the |Δω| values are reduced, indicating that the motions are generally smaller (Fig. 3D). Here, it is worth noting that we do not have coverage of the mini (A’)-helix and N-terminal end of the C-helix to determine whether they are also behaving distinctly as in S15: while we do observe conformational exchange in these regions of S1, the specific methyl probes undergoing exchange are different from those in the wild type protein, so cannot be directly compared. However, when we compare the two superkines, we observe a large cluster of well-correlated methyls with lower |Δω| values, and a cluster of methyls with much higher |Δω| values in S15 which comprise the A-helix and D-helix (Fig. 3E). Taken together, these results suggest that the superkine mutations alter dynamic transitions in the IL-2 hydrophobic core network, generating new and unique pathways by altering an intrinsic process that can be observed in all three proteins.

### IL-2 superkines show distinct conformational landscapes and allosteric networks in MD simulations

Given the interesting conformational exchange processes observed in our CPMG analyses, we reasoned that molecular dynamics (MD) simulations may uncover a clear picture of the underlying allosteric networks. MD simulations are a powerful tool for interpreting observed dynamics at a range of biologically relevant timescales that are accessible by NMR and other biophysical techniques, by delineating plausible allosteric pathways which connect functionally important sites in proteins^52^. To explore potential allostery within the proteins, we ran adaptive sampling simulations of wild type IL-2, S1, and S15. The adaptive sampling simulations were performed using a goal-oriented algorithm, fluctuation amplification of specific traits (FAST^53^) to maximize the root-mean-square deviation (RMSD) to the starting structure, allowing for more conformational sampling in a shorter amount of time. We analyzed the local structural transitions sampled in our MD simulations to explore agreement with our NMR experiments by monitoring alpha carbon root-mean-square fluctuation (RMSF) values and the sampling of different rotameric states gathered from chi2 dihedral angles (Fig. S11). We found that residues with increased structural variations, identified through high RMSF values and varied χ^2^ dihedral angle distributions, also show high R_ex_ values in our CPMG experiments, indicating that the MD simulations can be used to provide a plausible mechanism for the underlying structural changes observed in our NMR experiments and provide more insight into allostery within the IL-2 variants.

Unsurprisingly, we found that the superkines explore different conformational landscapes compared to wild type IL-2 (Fig 4A). We used the dimension-reduction method time-lagged independent component analysis (tICA) to show the conformational landscapes explored. The input features used for this analysis were pairwise distances between the methyls in exchanging residues identified in our CPMG NMR experiments (Table S2). As expected from our experimental results, we found significantly different conformational landscapes for each protein: although all three have similar “major” state conformations (at approximately IC1 and IC2 values of 0), their “minor” states each sample distinct conformational spaces. The two main motions that separate the conformational landscapes of the wild type and superkines are depicted in the tICA landscape (Fig. 4A & S12). The first main motion is along independent component 1 (IC1), where we observe the B-helix shifting up towards solution, and the C-helix shifting up and inwards towards the core. Since the loop between these helices (BC-loop) is very short in S1 (6 amino acids) and heavily mutated, the helices move together, with the C-helix slightly displacing the B-helix when it moves in towards the core, shifting the B-helix upwards. This motion is prominent in S1 but is not observed to the same extent in wild type and S15, likely because the BC-loop is longer (8 amino acids) and less rigid. The second main motion (along IC2) is concentrated within the mini (A’)-helix, where we observe partial unfolding in S15 and a shift inward toward the protein core in S1. Notably, the mini-helix is located at the IL-2Rα binding interface on IL-2, and has been shown to undergo a distinct conformational change upon receptor binding^39^. To better understand the structural differences between major and minor conformational states sampled by each protein, we extracted approximately 100 representative frames along IC1 and IC2 (Fig. S12, Video S1-12). In wild type IL-2, there is a pivot about the N-terminal end of the mini-helix, with the C-terminal end pushed out into solution in the minor state, which also causes a slight repositioning of the other helices. In S15, the mini-helix unfolds almost completely in the minor state, and the A- and D-helices lift up toward the protein core while the B-helix bends downwards and inwards, and the C-helix shifts downwards. In S1, the mini-helix remains well-folded and shifts inwards toward the protein core, causing the C-terminal end of the B-helix to pivot slightly toward the C-helix. Notably, the C-helix appears to be considerably more shifted in the S1 minor state compared to wild type IL-2, which only experiences a very minor shift at the N-terminal end of the C-helix. Given previous observations of C-helix conformational “priming” at the IL-2Rβ interaction site^7,21,22,37,39^, it is plausible that the conformational dynamics of the superkines enhance IL-2Rβ binding through a hydrophobic core repacking mechanism. Taken together, our tICA results support a model in which both superkine minor states achieve conformations that reposition the IL-2Rβ binding interface, which are not reached by wild type IL-2, via structurally distinct core repacking mechanisms largely enabled by distant motions at the mini-helix and B-helix.

**Figure 4:**
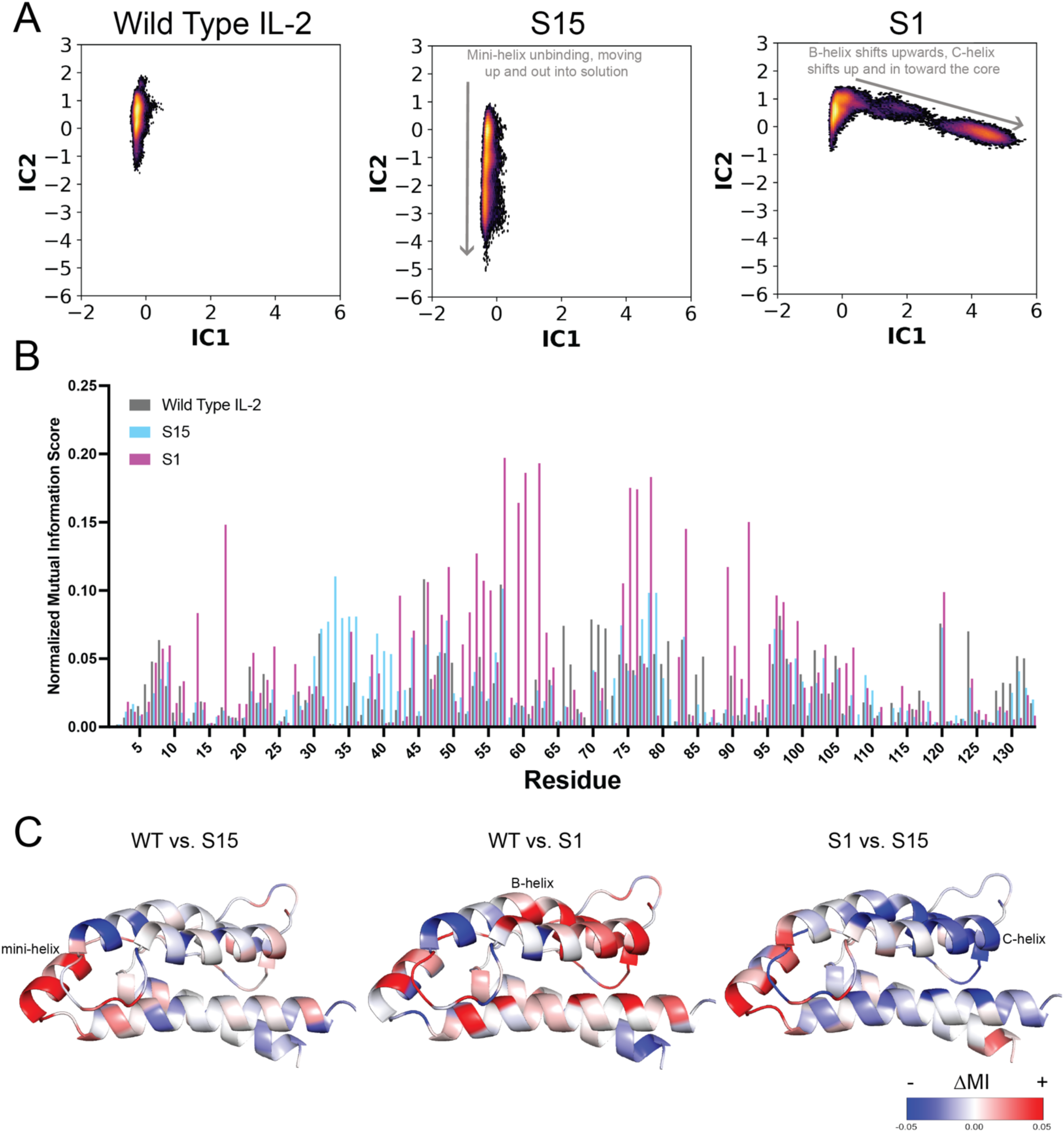
Molecular dynamics simulations show that superkines undergo distinctive core repacking mechanisms. Simulations were performed using RMSD-based adaptive sampling for a total of 5 μs of simulation time per protein. A) Extracted components of combined time-lagged independent component analyses for wild type IL-2 (left), S15 (center), and S1 (right) with a gradient portraying density of simulation frames ranging from low (purple) to high (yellow) density. B) CARDS mutual information scores for each residue in the wild type IL-2, S15, and S1, normalized based on the total number of residues in each protein. S1 residue numbering is based on alignment to the wild type sequence. C) Difference in mutual information scores (ΔMI) plotted on wild type IL-2 (left, PDB ID: 1m47), S15 (middle, PDB ID: 7raa), and S1 (right, PDB ID: 7ra9) structures as a gradient with an increase in MI shown in red and decrease shown in blue.

Since the tICA results showed different conformational landscapes, we hypothesized that three proteins use different allosteric networks. To quantify correlations within each protein structure derived in our MD simulation trajectories, thereby mapping any plausible allosteric networks, we employed Correlation of All Rotameric and Dynamical States (CARDS)^54^. We used CARDS mutual information (MI) data to identify key residues within each network (Fig. 4B). Mutual information scores define the strength of the correlation between two different residues in a protein, where a high value suggests a strong communication between those residues. Thus, the normalized sum of all mutual information scores for a chosen residue reveals the extent of correlation between that residue and all other residues in the protein, revealing the most important “control sites” for the overall allosteric network. Within wild type IL-2, we found the highest correlations within and between two clusters which represent the N-terminal end of the B-helix and parts of the hydrophobic core. Notably, in both S15 and S1 we observe clusters with increased MI scores relative to wild type IL-2 (Fig. 4C). In S15, the most prominent correlation is within the cluster representing the mini-helix (resid. 33-44). This mini-helix cluster is highly correlated with several residues at the N-terminal end of the B-helix and in the BC-loop near the IL-2Rβ binding region. This finding is consistent with our tICA results, where we found that S15 undergoes unfolding in the mini-helix, bending in the B-helix, and a resulting shift in the C-helix. S15 has nearby superkine mutations in the A’-loop, which may be responsible for destabilizing the mini-helix and causing unfolding in the minor state, leading to enhanced correlations along this region. On the other hand, S1 shows very high MI scores at sites within part of the A’B-loop and the N-terminal end of the B helix (resid. 47-64). We see significant correlation between this cluster and several other regions of the protein, including an extremely high correlation with the C-helix (Fig. S13). Notably, residues within the highly mutated BC-loop also have enhanced MI scores, indicating that the significant redesign of the BC-loop in this superkine likely contributes critical allosteric effects to the altered S1 network. Taken together, these results indicate that the mini-helix and B-helix regions are the most distinguishing clusters in the MD-derived allosteric networks of S15 and S1, respectively.

### Rational mutation of a key site in the hydrophobic core network disrupts superkine dynamics

Our combined NMR and MD analyses highlight differences in dynamics between wild type IL-2 and the two superkines, S15 and S1, uncovering the underlying allosteric networks. We hypothesized that the changes in the superkine core hydrophobic networks as a result of the mutations consequently contribute to the enhanced receptor binding which is characteristic of the superkine class of IL-2 mutants^21,22,55^. Thus, we rationally designed a point mutation at a key residue in the allosteric network of S1, with the goal of quenching the conformational exchange process. More specifically, we focused on the residues within the most correlated clusters in our CARDS datasets as well as sites that are involved in the conformational exchange process in our NMR data. We found that several residues with high mutual information values are all clustered near each other in the A’B-loop and B-helix (Fig. S14), and some of the same residues are undergoing conformational exchange based on CPMG relaxation dispersion profiles. We identified a key leucine residue, Leu56, which is buried in the S1 hydrophobic core beneath the B-helix, is central to the important region in the S1 allosteric network, and is not in proximity to any of the receptor binding sites. We hypothesized that a mutation at this site would disrupt the S1 allosteric network, so we explored this by performing adaptive sampling (FAST-RMSD) MD simulations to test this hypothesis. Our simulation for the S1 L56A mutant showed a clear disruption to the allosteric network (Fig 5A, S14).

**Figure 5:**
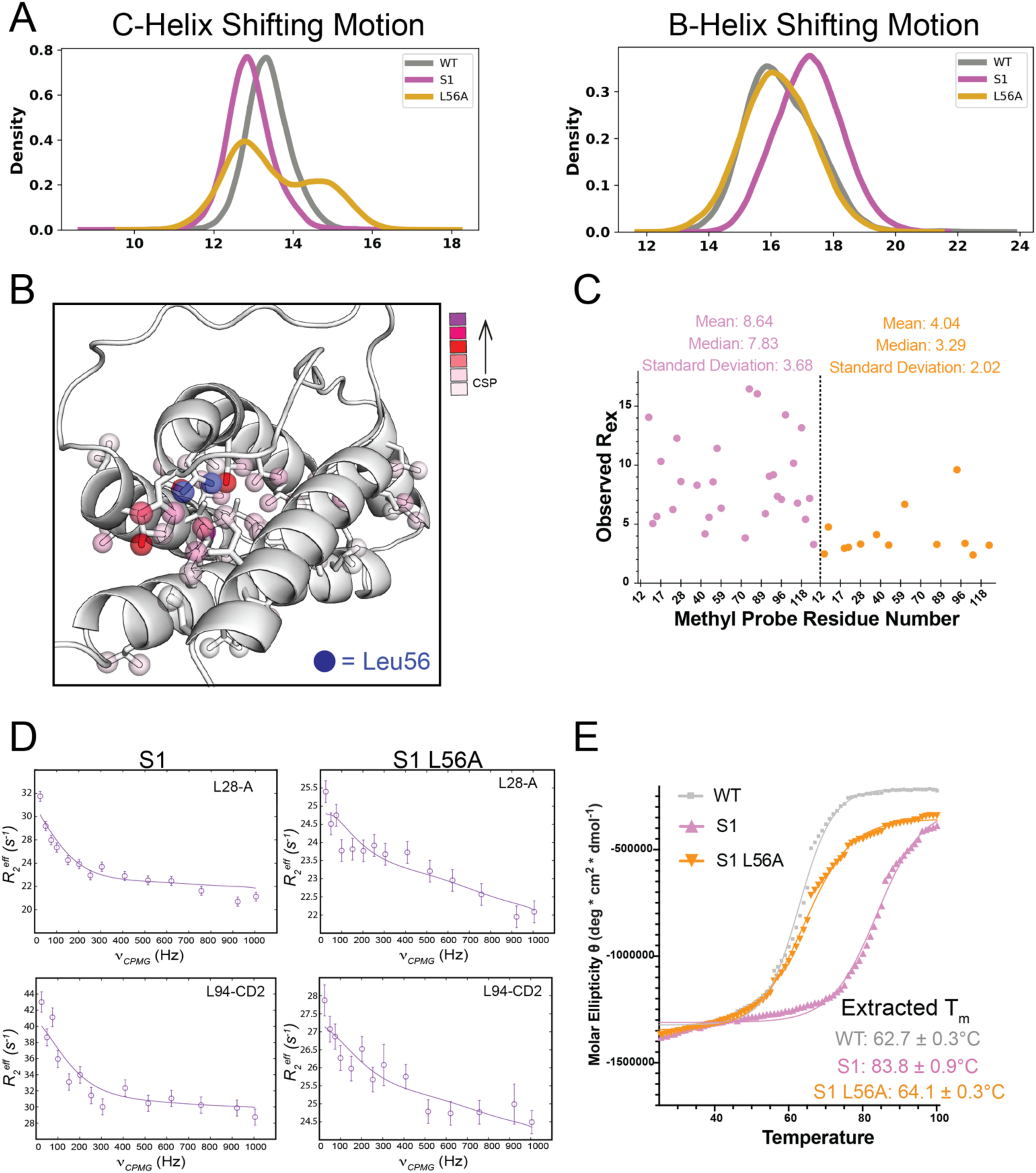
The L56A mutation disrupts the S1 allosteric network. A) Pairwise distances between residues with distinct motions capturing the behaviors of each protein. For the B-helix shifting distribution, the following pairwise distances were averaged: L12-L70, L18-L66, L14-L59, L21-L70. For the C-helix shifting distribution, the following pairwise distances were averaged: L18-L96, L18-V91, L17-V91, L14-V91. B) Methyl chemical shift perturbations quantifying spectral differences between S1 and S1 L56A plotted as spheres in a color gradient on the S1 crystal structure (PDB ID: 7ra9). The methyls at the L56A mutation site are plotted as blue spheres for reference. C) Rex profiles for S1 (pink) compared to S1 L56A (orange), where Rex is defined as R2^eff^ (25 Hz) – R2^eff^ (1005 Hz) from methyl-selective, single-quantum CPMG relaxation dispersion experiments collected at 800 MHz ^1^H magnetic fields at 5°C. Mean, median, and standard deviation are shown for reference. D) Representative dispersion curves extracted for S1 and S1 L56A, collected at 800 MHz ^1^H magnetic field. Shown are dispersion curves for Leu28-A (not stereospecifically assigned) and Leu94-CD2. Experimental data are shown as circles in all panels, with errors estimated from the S/N in the raw experimental data. The best-fit lines are shown for a global analysis using a two-site conformational exchange model of all methyls with relaxation dispersion profiles showing Rex > 3. E) Circular dichroism thermal melt curves collected at 222 nm in 1°C intervals from 25°C to 98°C for wild type IL-2 (gray), S1 (pink), and S1 L56A (orange) with extracted Tm values fitted from a nonlinear Boltzmann sigmoidal curve fit in GraphPad Prism.

Next, we expressed the S1 L56A mutant recombinantly to characterize its dynamics in solution (Fig. S1 & S15). At 25°C, we observe by circular dichroism that the overall secondary structures of wild type IL-2, S1 superkine, and S1 L56A are very similar (Fig. S15E). We assigned the methyl side chains of selectively ^13^C/^1^H-labeled isoleucine, leucine, and valine residues on the S1 L56A mutant and calculated CSPs relative to the S1 superkine (Figure 5B, S15B-D). While most of the large CSPs are found at methyl probes near Leu56 in the crystal structure, we also observe smaller CSPs at distal sites, suggesting that the L56A mutation has allosteric effects throughout the entire S1 core. Next, we recorded CPMG relaxation dispersion experiments at 800 MHz ^1^H magnetic field to verify that the L56A mutation alters S1 dynamics in solution. We observed that the S1 L56A mutant has a reduced R_ex_ landscape compared to the S1 superkine (Fig. 5C). While we did not pursue a quantitative global fit of this data with a two-site exchange model, a close inspection of the dispersion curves allows us to observe that the rate of decrease in R_2_^eff^ as a function of CPMG pulse frequency is very different when comparing curves from S1 and S1 L56A, suggesting that the exchange rate (k_ex_) of the conformational exchange process is likely faster (Fig. 5D). These results demonstrate that we can rationally perturb the S1 allosteric network by introducing a point mutation guided by our NMR and MD simulation analysis.

However, we note that the thermal stability of the S1 L56A mutant is reduced compared to S1, which was engineered to be highly stable, resulting in a 20°C increase in T_m_ compared to wild type IL-2 (Fig. 5E)^22^. Given several previously reported examples highlighting an inverse correlation between protein conformational exchange kinetics and thermal stability^56–60^, it is not surprising that a disruption of S1 dynamics could reduce the protein stability at high temperatures. Nevertheless, to verify whether a reduction in thermal stability alone is sufficient for disrupting IL-2 dynamics and receptor interactions, we explored an additional S1 mutation, V84A. Residue 84 is an S1 mutation site (mutated from an Ile in wild type IL-2 to a Val in S1) which was included in the S1 design with the goal of improving core packing, thereby enhancing the thermal stability of the molecule^22^. By mutating this residue – which is not predicted to be involved in the dynamical or allosteric processes defined by our NMR data and adaptive sampling MD simulations – to alanine, we reduced S1 thermal stability without impacting S1 dynamics, allosteric networks, or receptor binding (Fig. S16). Taken together, these results suggest that reducing S1 stability alone does not necessarily disrupt IL-2 allostery or receptor binding, as demonstrated by the V84A mutation. This further supports the critical role of the core allosteric network identified by our combined NMR/MD studies for regulating interactions with co-receptors.

### Perturbing the S1 core network partially reverts receptor binding and signaling function

It has been previously suggested that the mechanism for enhanced superkine interactions with IL-2Rβ involves stabilization of the C-helix, promoting a primed conformation for receptor recognition^21,22^. We further hypothesized that superkine conformational switching occurs via a more complex core repacking process required for enhanced receptor binding. To test this, we measured binding to IL-2Rβ by surface plasmon resonance (SPR). As previously described, we observed that the S1 superkine showed significantly enhanced binding affinity, with an approximately 20-fold reduced equilibrium dissociation constant (K_D_) relative to WT IL-2 at approximately 18.60 ± 2.59 and 385.1 ± 11.3 nM, respectively (Fig. 6A-B)^22^. Due to the very fast association and dissociation rates, we could not fit the wild type IL-2 kinetics. However, examination of the binding curves reveals that S1 achieves a slower dissociation rate constant (0.0184 ± 0.0020 s^-1^), thereby prolonging the average lifetime of the complexed state. In comparison, we observed that the S1 L56A mutant had more than two-fold reduced binding affinity compared with S1, with a K_D_ of 46.2 ± 4.82 nM, driven by a twofold increase in the dissociation rate constant (0.0393 ± 0.0070 s^-1^, Fig. 6C). Thus, the L56A mutation on S1, which disrupts the S1 dynamics, causes an increase in the dissociation rate constant compared to S1, confirming that conformational dynamics can impact the average lifetime of the receptor-bound state.

**Figure 6:**
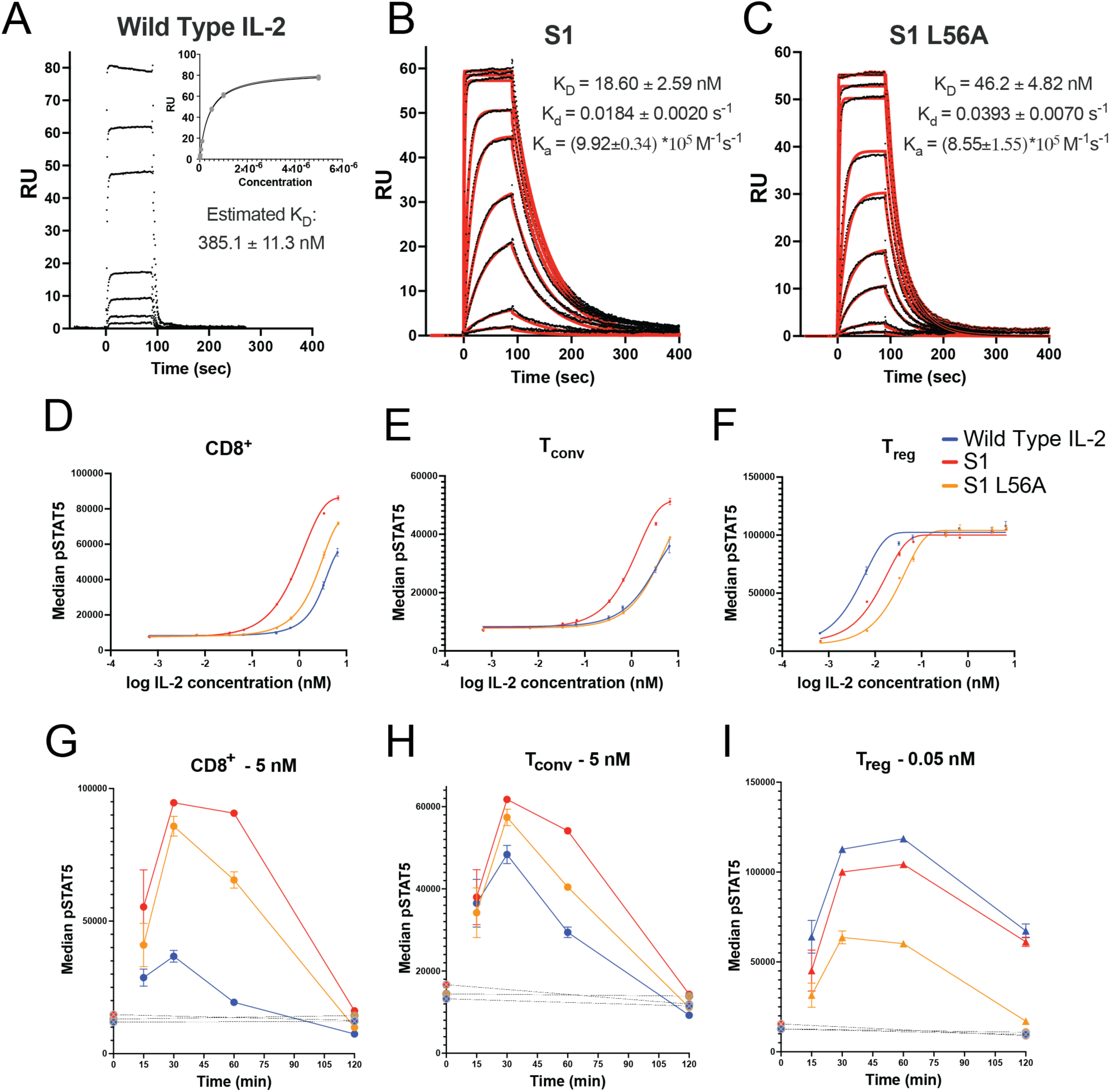
Dysregulation of S1 dynamics disrupts receptor binding. Steady-state binding affinity of A) wild type IL-2, B) S1, and C) S1 L56A mutant against IL-2Rβ, measured by surface plasmon resonance. Wild type IL-2 could not be fitted kinetically within the limitations of our instrument, so the raw binding curve and affinity fit used to extract the KD is shown. For S1 and S1 L56A, red fitted curves represent kinetic best fits to the data, and all extracted kinetic parameters and KD values are shown. Pre-isolated T cells from human donors and incubated with IL-2 variant for 20 min at 37°C. Dose-response curves monitoring median fluorescence intensity of STAT5 phosphorylation are shown in in D) CD8^+^ T cells, E) CD4^+^ Tconv cells, and F) CD4^+^ FoxP3^+^ Treg cells. G-I) IL-2 pulse experiments covering longer time courses ranging from 15 to 120 minutes at 37°C. In these experiments, pre-isolated T cells were first chilled and coated with IL-2 variant on ice, washed, and treated with an anti-IL-2 blocking antibody at the start of the 37°C incubation to prevent autocrine IL-2 production during the experiment. Control samples chilled and treated with IL-2 but not anti-IL-2 blocking antibody and kept on ice during the time course are shown as baseline controls (marked by dotted lines). Dose-response curves monitoring median fluorescence intensity of STAT5 phosphorylation are shown in *G*) CD8^+^ T cells treated with 5 nM of IL-2 variant, H) CD4^+^ Tconv cells treated with 5 nM of IL-2 variant, and I) CD4^+^ FoxP3^+^ Treg cells treated with 0.05 nM of IL-2 variant.

Finally, to connect our biophysical findings with IL-2 signaling function, we tested the impact of the L56A mutation on activation of primary T cells from human donors. We monitored phosphorylation of signal transducer and activator of transcription 5 (STAT5) in response to each IL-2 variant by phospho-flow cytometry^61,62^. To determine the functional activity of the S1 L56A mutant, we probed its effects on IL-2-induced signaling in CD8^+^ T, CD4^+^ conventional T (T_conv_), and CD4^+^ FoxP3^+^ T_reg_ cell subsets (Fig. 6D-F, S17-18). Consistent with our SPR results, S1 L56A showed a reduction in phosphorylation of STAT5 compared to S1, but still maintained enhanced signaling compared to wild type IL-2 in CD8^+^ T cells (Fig 6D) and maintained similar levels of signaling as wild type IL-2 in T_conv_ cells (Fig 6E). Using CD8^+^ T cells, which primarily express the dimeric (IL-2Rβγ_c_) receptor under resting conditions (Fig. S18), we consistently observe an approximately 2.5x increase in EC_50_ value for S1 L56A compared to S1 across multiple experiments, and an approximately 10x increase in EC_50_ value for wild type IL-2 compared to S1 (Fig. S17). These results are consistent with previous analyses which demonstrated that disrupting IL-2Rβγ_c_ binding enhances T_reg_ bias over CD8^+^ T and T_conv_ cells (Fig. S17)^33^. Notably, S1 was designed to function independently of IL-2Rα with an included mutation to inhibit binding^22^, thus it was unsurprising that both S1 and S1 L56A had reduced T_reg_ bias compared to wild type IL-2 since T_reg_ cells express IL-2Rα at high levels (Fig. S17). Given that autocrine production of IL-2 is possible especially in antigen-experienced T cells^63–65^, we also performed a pulse-chase experiment in which pre-chilled cells were treated with IL-2 variant on ice for 30 minutes, washed, and then treated with an anti-IL-2 blocking antibody prior to a time-course incubation at 37°C (Fig. 6G-I). Our results show a similar trend, in which S1 L56A has diminished signaling compared to S1 in both CD8^+^ T cells and T_conv_ cells, while still maintaining improved signaling compared to wild type IL-2 (Fig. 6G-H). Consistently, in T_reg_ cells we observe reduced signaling of both S1 and S1 L56A compared to wild type IL-2, as expected (Fig. 6I). In summary, these results reveal that altered core allosteric pathways in superkines contribute to enhanced receptor binding, while disruption of these pathways diminishes signaling function *in vitro*.

## Discussion

Several groups have developed IL-2 variants with enhanced T cell signaling properties by either favoring or disfavoring interactions with specific receptor subunits. Designed IL-2 mutations for cancer therapy aim to increase the affinity of IL-2 for the IL-2Rβγ_c_ receptor expressed on T_eff_ cells, and/or to disrupt the IL-2Rα binding interface to reduce effects on T_reg_ and other IL-2Rα^+^ cells for cancer therapies^21–27^. On the other hand, IL-2 modifications designed for autoimmune therapies aim to preferentially induce signaling through the trimeric high-affinity receptor (IL-2Rαβγ_c_), stimulating T_reg_ cells with greater specificity and thereby widening the clinical dosing window^28–32,66^. Aside from the superkines, previous clinical translation of IL-2 mutants has principally relied on modulating IL-2 residues that directly interact with receptor subunits, which typically causes a strong loss in signaling potency and thereby presents a barrier to achieving greater selectivity toward desired T cell subsets^33^. As an alternative, finetuning IL-2’s core allosteric network to design an orthogonal set of IL-2 functional mutants has not been explored in depth, given that the underlying molecular mechanisms had not previously been investigated. The current study provides a foundation for future design efforts by identifying the residues involved in the cytokine’s core hydrophobic network that could be exploited for engineering novel functional agonists with enhanced properties.

Prior to this study, there have been a limited number of investigations into IL-2 allostery. In crystal structures, it had been observed that “priming” of IL-2 by binding of IL-2Rα causes a subtle conformational change at the IL-2Rβ binding interface on the opposite side of the protein, which was believed to enhance the binding of this subunit to the receptor complex^7,37,39^. This has been corroborated by extensive binding studies which demonstrate that pre-binding of IL-2Rα increases the affinity of IL-2 for IL-2Rβ^38^. Furthermore, computational approaches have previously been employed to understand the correlations between individual superkine mutation sites or between small molecule binders and other residues in the protein: in agreement with our results presented here, correlations were observed between residues on opposite sides of IL-2, including residues in the BC-loop, B-helix, C-helix, and mini (A’)-helix^67,68^. Our study represents the first direct experimental investigation into IL-2 dynamics in solution and provides a more complete understanding of how protein dynamics affect receptor interactions. Furthermore, the key differences in core dynamics between wild type IL-2 and the two superkines are difficult to access by other experimental methods, highlighting the power of solution NMR for parsing these features. By combining experimental and computational approaches, we have provided a detailed view of the core repacking events at play in each protein and identified the specific network of residues involved in each process.

Previous studies have proposed that stabilizing the IL-2 C-helix in a “primed” conformation promotes higher affinity receptor binding, and the superkines S15 and S1 were designed to achieve this through stabilizing the N-terminal loop ahead of the C-helix (BC-loop) and optimizing core packing nearby^7,21,22,37–39^. Thus, we expected that the dynamics of the superkines would be partially quenched relative to wild type IL-2, especially at regions near the C-helix. Instead, we observed that the solution dynamics of these superkines are very different from those of wild type IL-2 but are certainly not quenched on the microsecond to millisecond timescale by NMR. In fact, in the case of S15 it appears that dynamics are enhanced compared to wild type IL-2, as suggested by a cluster of residues with higher magnitudes of conformational exchange (R_ex_) and higher |Δω| values indicating larger motions resulting from the switch between the major and minor states (Fig. 3D, S6). In the case of S1, we observe relatively similar R_ex_ profiles to wild type IL-2 overall, but considerably slower conformational exchange kinetics. We also note that substantially more methyl probes are participating in each superkine dynamical process than observed in wild type IL-2. Thus, we propose that the mechanisms by which the S15 and S1 superkines achieve higher affinity binding to IL-2Rβ are more complex than has been suggested previously, and likely involve a combination of features which are distinctive to each superkine. Our data suggest that a hydrophobic core repacking mechanism is at play in wild type IL-2, which is altered by the superkine mutations to generate new allosteric pathways that are both structurally distinct and kinetically slower than the wild type conformational exchange process.

We rationally designed an S1 mutant, L56A, to demonstrate that mutations disrupting superkine dynamics impact receptor binding and signaling (Fig. 6). Interestingly, |Δω| values for Leu56 are consistently well correlated across wild type IL-2, S15, and S1 in our CPMG data (Fig. 3C-E), and Leu56 serves as a linchpin in the allosteric network observed in our MD simulations (Fig. 4). Furthermore, it has been shown that mutations at Leu56 on wild type IL-2 are highly detrimental to IL-2R interactions, causing a severe loss in binding capabilities^69^. We therefore suggest that our L56A mutation on S1 disrupts intrinsic core dynamics shared by all three IL-2 variants, like a molecular “off switch” to reduce the correlations within the network. It is likely that the binding and function of S1 L56A is partially rescued by the superkine mutations, which retain some dampened dynamical properties of S1, as observed in our NMR data (Fig. 5C-D). While we observed that S1 L56A has lower thermal stability than the highly stable S1 superkine, we also qualitatively observed that S1 L56A dynamics are likely faster than the process observed in S1. We propose that the original design principles for S1, which aimed to hyper-stabilize IL-2 to achieve higher binding affinity^22^, inadvertently selected for an optimized allosteric network with a slower conformational exchange process. Thus, a mutation intended to disrupt this network would both increase the conformational exchange rate and reduce receptor binding affinity via an increase in the dissociation rate constant, as observed in our SPR binding studies (Fig. 6B-C). We therefore postulate that core dynamics, thermal stability, and binding affinity may be intertwined in this system, where mutations that increase the rate of conformational exchange (k_ex_) also decrease the binding affinity for IL-2Rβ by increasing the dissociation rate constant. However, we also demonstrated that a mutation that disrupts S1 thermal stability, but is not predicted to disrupt the allosteric process, does not significantly impact function in T cells (Fig. S16). Thus, core dynamics are key to predicting non-receptor interface mutations that will modulate signaling function, rather than thermal stability alone.

From a practical perspective, the T_reg_-biasing applications in autoimmune disease treatment serve as a pertinent example for such design endeavors. Existing T_reg_-biasing strategies either *i)* increase the affinity of IL-2 for IL-2Rα or *ii)* decrease the affinity for IL-2Rβγ, or both, by altering the receptor binding interfaces on IL-2^24,29–32,34^. These approaches rely heavily on differentials in IL-2Rα expression across T cell subsets (which vary between patients), limiting the capacity to distinguish between immune cell subsets for specific activation of T_reg_ cells. As an alternative approach, our work highlights the allosteric pathways that connect the IL-2Rα and IL-2Rβ binding surfaces through IL-2’s hydrophobic core, which could be exploited to fine-tune binding cooperativity. Future work will aim to identify rational mutations designed to alter protein dynamics, toward enhanced cooperativity between IL-2Rα and IL-2Rβ, making IL-2 binding more favorable in the presence of IL-2Rα and achieving greater T_reg_ selectivity. In summary, our work has identified a mechanistic role for IL-2 core dynamics in receptor binding and T cell signaling function, which are altered by the superkine mutations to achieve distinctive, functionally enhanced binding and signaling properties. These results provide a basis for further exploration of the allosteric networks of IL-2 and other common gamma-chain cytokines.

## Materials and Methods

### IL-2 expression, refolding, and purification

Codon-optimized DNA encoding human IL-2 with a site-specific mutation (C125S) was expressed in BL21 (DE3) *E. coli* cells into inclusion bodies by induction with 1 mM IPTG at OD_600_ = 0.6 followed by cell incubation at 37°C for 5 hours or at 22°C for 18 hours, at 200 RPM. After solubilizing the purified inclusion bodies in guanidine-HCl, ∼30-60 mg of inclusion bodies were rapidly diluted dropwise into 200 mL of refolding buffer (1.1 M guanidine, 6.5 mM cysteamine, 0.65 mM cystamine, 110 mM Tris, pH 8.0) at 4 °C while stirring. Refolding proceeded overnight at 4 °C without stirring. The solution was dialyzed into a buffer of 20 mM MES (pH 6.0), 25 mM sodium chloride. Purification of refolded IL-2 was performed by SEC with a HiLoad Superdex 75 16/600 column (Cytiva) at 1 mL/min with running buffer (50 mM NaCl, 20 mM sodium phosphate pH 6.0).

### Expression and purification of IL-2 receptor beta chain

The human IL-2Rβ ectodomain gene (amino acids 1–214) containing a C-terminal biotin acceptor peptide and hexahistidine tag, was cloned into the gWiz vector (Genlantis) via isothermal assembly. The resulting plasmid was then validated by Sanger sequencing. The human IL-2Rβ was then expressed recombinantly in human embryonic kidney (HEK) Expi293 cells. The HEK Expi293 cells were grown to 1.0-1.5×10^6^ cells/mL on the day of transfection. The plasmid DNA and polyethyleneimine max (PEI max, Polysciences) were separately diluted to 1μg/mL and 0.05 mg/mL, respectively, in Opti-MEM medium (Gibco) and incubated at room temperature for 10 min. Subsequently, the DNA/PEI max mixture was added to a flask containing the HEK Expi293 and then incubated at 37°C with shaking for 5 days. The secreted protein was then harvested from HEK Expi293 cell supernatants by Ni-NTA (Thermo Scientific) affinity chromatography. The extracted human IL-2Rβ protein was then biotinylated with the soluble BirA ligase enzyme in 0.05 M Bicine pH 8.3, 10 mM ATP, 10 mM magnesium acetate, and 50 μM biotin (Avidity) for 1 hour at room temperature, followed by an overnight incubation at 4°C. Excess biotin was then removed by size-exclusion chromatography on an ÄKTA^™^ FPLC instrument using a Superdex 200 column (Cytiva). The protein biotinylation and purity were then verified with a streptavidin gel-shift assay via SDS-PAGE analysis.

### Preparation of NMR samples

All proteins were overexpressed in M9 medium supplemented with the appropriate precursors for the desired labeling scheme as described in detail previously^40,70^. Methyl labeling on a triple-labeled background, referred to as **ILV\***, was achieved by the addition of methyl-selective precursors Ile-δ_1_-[^13^CH_3_], Leu-δ-[^13^CH_3_/^12^C^2^H_3_], Val-γ-[^13^CH_3_/^12^C^2^H_3_] (Sigma-Aldrich) in an M9 media containing ^2^H_2_O, ^2^H,^13^C glucose (Sigma #552151), and 1 g/L of ^15^NH_4_Cl. So-called **deuterated ILV-methyl** (Ile ^13^Cδ_1_; Leu ^13^Cδ_1_/^13^Cδ_2_; Val ^13^Cγ_1_/^13^Cγ_2_) U-[^15^N, ^2^H]-labeled proteins were prepared in M9 medium in ^2^H_2_O, supplemented with 2 g/L of ^2^H,^12^C glucose (Sigma #552003) and 1 g/L of ^15^NH_4_Cl. Similarly, **non-deuterated ILV-methyl** (Ile ^13^Cδ_1_; Leu ^13^Cδ_1_/^13^Cδ_2_; Val ^13^Cγ_1_/^13^Cγ_2_) U-[^15^N]-labeled proteins were prepared in M9 medium in water, supplemented with 1 g/L of ^15^NH_4_Cl. **U-[^15^N]-labeled** proteins were expressed in M9 medium in water containing 1 g/L of ^15^NH_4_Cl only. Samples in the concentration range of 50 to 300 μM were prepared in a standard NMR buffer (50 mM NaCl, 20 mM sodium phosphate pH 6.0, 0.001 M sodium azide, 5% ^2^H_2_O).

### Backbone and methyl NMR assignments

Wild type IL-2 assignments have been published previously (BMRB 28104)^41^. Following similar methods, the S1 and S15 superkines were assigned as follows: We obtained backbone assignments using TROSY-based 3D HNCA, HN(CA)CB, and HNCO experiments^71–74^ recorded with ILV* samples. Backbone amide assignments were confirmed using amide-to-amide NOEs obtained from 3D Hall-NHn SOFAST NOESY-HMQC experiments^75^. Final backbone assignments were further validated using TALOS-N^42^. Next, Ile, Leu, and Val methyl assignments were achieved using methyl-to-methyl NOEs observed in 3D Cm-CmHm SOFAST NOESY-HMQC experiments (mixing times: 50, 150, 300 msec) in addition to methyl-to-amide NOEs observed in 3D HnHa-CmHm SOFAST NOESY-HMQC experiments^75^ (mixing times: 150 msec) recorded on deuterated ILV methyl-labeled samples^70,76^. Leu/Val geminal pairs were determined by comparing NOE strips in 3D Cm-CmHm SOFAST NOESY experiments recorded using short (50 ms) and long (300 ms) mixing times. Ile δ_1_ methyl types were identified by their characteristic upfield chemical shifts. Leu and Val methyl types were identified using phase-sensitive 2D ^1^H-^13^C HMQC experiments recorded on the ILV*-labeled sample^77,78^. We did not pursue stereospecific methyl assignments for the S1 and S15 Leu and Val assignments; instead, geminal pairs that we did not stereospecifically assign were arbitrarily marked as “A” and “B”. To validate ambiguous methyl assignments, we employed site-directed mutagenesis of L25 and L70 in S15 and L25 and V89 in S1 and prepared non-deuterated ILV samples to collect 2D methyl HMQC experiments. S1 L56A methyl assignments were transferred from the S1 assignments using 3D Cm-CmHm NOESY experiments. For chemical shift perturbation calculations, ^15^N TROSY HSQC or methyl 2D SOFAST ^1^H-^13^C HMQC NMR spectra were collected at 800 MHz ^1^H magnetic fields. Amide backbone chemical shift perturbations were calculated using the following equation, given the ^15^N and ^1^H chemical shifts: 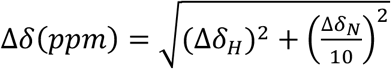. Similarly, methyl chemical shift perturbations were calculated using the following equation, given the ^13^C and ^1^H chemical shifts: 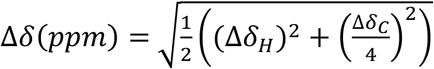. All the data were recorded at a ^1^H magnetic field of 600 or 800 MHz at 25°C. All spectra were acquired using Topspin 4 acquisition software from Bruker. The data were processed in NMRPipe^79^ and analyzed in NMRFAM-SPARKY^80^ and POKY^81^ programs.

### Nuclear spin relaxation experiments

We utilized pulse sequences for the measurement of ^15^N T_1_, T_2_, and ^1^H-^15^N heteronuclear NOE as described previously^82–85^. All experimental data were recorded at 600 and 800 MHz ^1^H magnetic fields on U-[^15^N]-labeled NMR samples at 200-300 μM concentrations. T_1_ measurements were performed using a series of 12 experiments, with relaxation delays ranging from 20 to 1000 msec at both magnetic fields. T_2_ measurements were performed using a series of 12 experiments, with relaxation delays within the range 16.96–271.36 msec at 600 MHz and within the range 16.96–237.44 msec at 800 MHz. Heteronuclear NOE spectra were collected with and without proton saturation at both fields. Peak intensities were extracted using POKY^81^ and subsequently used to obtain an estimate of the relaxation rates R_1_ and R_2_ by fitting to a two-parameter exponential decay model. Uncertainties in the fitted parameters were estimated by calculating error in peak heights from repeated measurements of one relaxation delay across the T_1_ and T_2_ series three times (60 msec for T_1_; 33.92 msec for T_2_). Error in heteronuclear NOE was propagated from the signal-to-noise of the reference and attenuated spectra. Optimization of the diffusion tensor, calculation of the overall molecular diffusion (tm), and model-free analysis of the relaxation data was performed with the program Relax^46–49^ using the fully automated protocol and default inputs. After the analyses were completed, R_1_, R_2_, and heteronuclear NOE values were back-calculated at 600 and 800 MHz to verify agreement between the model fitting and the experimental data.

### Methyl CPMG relaxation dispersion experiments

Methyl single-quantum ^13^C CPMG relaxation dispersion experiments^50^ were recorded on deuterated ILV-methyl labeled WT, S1, and S15 IL-2 samples (300 μM protein concentration) at ^1^H magnetic fields of 600 MHz and 800 MHz at 5 °C, on Bruker spectrometers equipped with a cryoprobe. The CPMG datasets were acquired as pseudo 3D experiments with a constant relaxation time period T_relax_ of 40 ms and with 15 CPMG pulse frequencies ν_CPMG_ = 1/(2τ) ranging from 25 to 1000 Hz, where τ is the delay between the consecutive 180° refocusing pulses in ^13^C CPMG pulse train. Relaxation dispersion profiles were calculated from peak intensities recorded at different CPMG frequencies using the following equation: R_2,eff_(ν_CPMG_) = -1/T_relax_ ln(I/I_0_), where I is signal intensity in the spectra collected at T_relax_ = 40 ms, I_0_ is signal intensity in the reference spectrum recorded at T_relax_ = 0. All data were processed using NMRpipe^79^ and peak intensities were picked using POKY^81^. The error was propagated from the noise level of the spectra. The variation in R_2,eff_ with ν_CPMG_ was fit to a two-site model of conformational exchange based on the Bloch-McConnell equations to extract values of exchange parameters (p_B_, k_ex_=k_AB_+k_BA_), as well as ^13^C chemical shift differences (|Δω|) for nuclei interconverting between states, using the software CATIA^51,86^. Final global fits for each of the three proteins included all methyl probes with R_ex_ (defined by the difference in R_2,eff_ at ν_CPMG_ = 25 and 1000 Hz) ≥ 3 at 800 MHz 1H magnetic field with acceptable fits across two magnetic fields in the two-site exchange model (9 methyl probes for wild type, 20 probes for S15, 19 probes for S1). The fitting via CATIA was performed by minimizing the function chi^2^ as previously described^40,87^. Chi^2^ surface plots were generated to evaluate the robustness of the extracted exchange parameters (p_B_, k_ex_) in the context of the global fits. Numerical fitting was performed using the CATIA program as before, with pb and kex sampled from a grid with values ranging from 0-30% and 0-4000 s^-1^, respectively, where |Δω| was free to change during the chi^2^ minimization procedure.

### Molecular dynamics simulations

Molecular dynamics simulations were seeded from x-ray crystallography structures using the GROMACS^88^ software (version 2023.2) and the CHARMM36m force field^89^. The proteins were each solvated using TIP3P^90^ waters and 0.1 uM NaCl in a dodecahedron box. Energy minimization was performed using steepest descent minimization, followed by NVT equilibration for 200 ps, NPT equilibration for 1 ns. FAST-RMSD^53^ simulations were performed with 10 generations with 10 runs of 50 ns each for a total of 5 μs of simulation time per protein. Time-lagged independent component analysis plots were generated from pairwise distances between CD1 or CG1 atoms of isoleucine, leucine, and valine residues undergoing conformational exchange in at least one of the three proteins (based on NMR CPMG relaxation dispersion data), omitting mutation sites. All input files, starting structures, and analysis files are deposited in the following GitHub repository: https://github.com/ssolieva/IL2-dynamics.

### Allosteric network analysis from MD simulations

To create an allosteric communication map across the protein, we calculated mutual information between rotameric states of dihedral angles using a method called Correlation of All Rotameric and Dynamical States (CARDS)^54^. Categorizing dihedrals angles from both the side chain and backbone into greater rotameric states allows us to track an angle’s motions across the length of the simulation. To account for disordered communication, we additionally assigned states into ordered or disordered states based on whether they are stably in a rotameric state or transitioning between states respectively using the formula 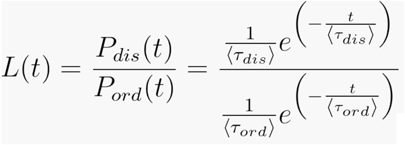. This formula is derivative from interpretations of Bayes factors, where P_dis_ is the probability of a state being disordered, Pord is the probability of a state being ordered, and their respective tau values represent mean ordered times. State determination is disordered if L > 3.0 and ordered otherwise. We then used mutual information to assess the level of correlated motions between these metrics using the formula: 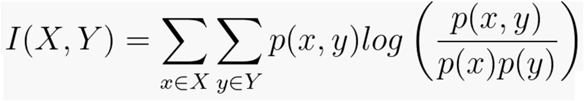, where x ∈ X refers to all the possible states (x) that a dihedral (X) can adopt, p(x) is the probability that the dihedral adopts the specific state x, with the same holding true for state y and dihedral Y. Thus, p(x,y) is the joint probability that dihedral X adopts state x and dihedral Y adopts state y. This mutual information is then normalized using the formula: 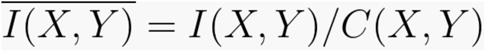, where C(X,Y) refers to the channel capacity which represents the maximum possible MI between two dihedrals. To capture the holistic correlation (IH), we can now use our normalized MI as such: 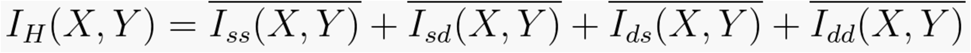, where Iss refers to the structure-structure correlation, Isd refers to the structure-disorder correlation, Ids refers to the disorder-structure correlation, and Idd refers to the disorder-disorder correlation. This allows for comparison between different types of correlations that can have different numbers of states. An example of this side-chain dihedrals, which have three possible rotameric states, so the maximum structural correlation (Iss) is log(3), but only two possible dynamical states so the maximum disordered correlation (Idd) is log(2). We identified a highly correlated set of residues by finding the residues that had the highest mutual information with our residues of interest.

### Site-directed mutagenesis

IL-2 plasmids were mutated by PCR amplification using primers containing the mutations of interest the KAPA HiFi HotStart ReadyMix PCR kit (Roche) according to the manufacturer’s recommendations. Template DNA was removed by DpnI digestion for 60 minutes at 37°C and transformed into *E. coli* DH5α strain and plasmids were isolated using the QIAprep Spin Miniprep Kit (QIAGEN). The introduced mutations and the absence of secondary mutations were verified by sequencing of plasmid DNA.

### Circular dichroism

CD spectra were measured on a JASCO 1500 CD instrument in a 1 cm pathlength cuvette. Protein samples were at ∼5 μM in 20 mM sodium phosphate (pH 6) and 50 mM NaCl. Thermal melt curves were obtained by monitoring the CD signal at 222 nm in 1 °C increments every 1 min from 25 to 95 °C, with 10 s of equilibration time, 8 s of digital integration time, and 3 mm bandwidth. Additionally, wavelength scans were collected in 10 °C increments across 190-260 nm with a continuous scan of 50 nm/min. All results were analyzed in GraphPad Prism v10 and melt curves were fitted to a nonlinear Boltzmann sigmoidal curve to estimate the T_m_.

### Liquid Chromatography – Mass Spectrometry

30 μM of protein were either untreated (oxidized) or treated with 2 mM DTT (reduced), incubated at room temperature for 10 minutes, and then diluted to a final concentration of 2 μM in a low pH buffer (20 mM sodium phosphate, pH 2.5, 50 mM NaCl). Samples were injected for LC-MS/MS in which integrated pepsin digestion was performed using a C8 5 μM column and a Q Exactive Orbitrap Mass Spectrometer. Peptide fragments corresponding to wild type IL-2, S1, or S1 L56A were identified using Thermo Proteome Discoverer v2.4. ExMs2 program was used to identify and analyze peptides.

### Surface plasmon resonance

BSP-tagged IL-2Rβ was biotinylated using the BirA biotin-protein ligase bulk reaction kit (Avidity) according to the manufacturer’s instructions, and biotinylation efficiency was evaluated by an SDS-PAGE gel shift assay. SPR experiments were performed in a BiaCore X100 instrument (Cytiva) in SPR buffer (150 mM NaCl, 20 mM sodium phosphate pH 7.4, 0.1% Tween-20) as described previously^91^. Approximately 300 resonance units (RU) of biotinylated IL-2Rβ were conjugated on a streptavidin-coated chip (Cytiva) at 10 μl/min. Various concentrations of IL-2 variant were flown over the chip for 60 seconds at 30 μl/min followed by a buffer wash with a 180 second dissociation time at 25°C. SPR sensorgrams, association/dissociation rate constants (k_a_, k_d_), and equilibrium dissociation constant K_D_ values were analyzed in BiaCore X100 evaluation software (Cytiva) using kinetic analysis settings of 1:1 binding. SPR sensorgrams and affinity-fitted curves were analyzed in GraphPad Prism v10.

### STAT5 signaling assays

All primary T cells used in this study were provided by the Human Immunology Core at the University of Pennsylvania under a protocol approved by the Children’s Hospital of Philadelphia Institutional Review Board. Approximately 0.1-0.5 million pre-isolated total T cells per well were resuspended in Advanced RPMI Complete medium (composed of Advanced RPMI (Thermo Fisher) supplemented with 10% heat-inactivated fetal bovine serum (FBS; Thermo Fisher), 1x penicillin and streptomycin solution (Millipore Sigma), 1x GlutaMAX (Thermo Fisher), and 10 mM HEPES (Thermo Fisher)) and plated into 96-well plate. For the continuous signaling experiments, cells were stimulated for 20 minutes at 37°C with serial dilutions of IL-2 (diluted in Advanced RPMI Complete medium) at a total volume of 100uL/well. For the IL-2 pulse experiment, ice cold cytokine at the desired concentration was added to pre-chilled cells and incubated for 30 min on ice. Cells were then washed three times with 1 mL of ice-cold Advanced RPMI Complete medium, spun at 500 x g for 5 min, and decanted. After the last wash, cells were resuspended in pre-warmed (37°C) media containing anti-cytokine blocking antibody (AB12-3G4, ThermoFisher Scientific 16-7027-85) at a final concentration of 20 ug/mL and incubated at 37°C for the following time points: 15, 30, 60, and 120 minutes.

After the appropriate treatment, cells were immediately fixed by addition of 1 mL of 1× TFP Fix/Perm buffer (Transcription Factor Phospho Buffer Set, BD Biosciences) and incubated at 4°C for 50 minutes on ice. 350 μL 1× TFP Perm/Wash buffer (BD Biosciences) was then added to each well, and cells were pelleted and washed again with 750 μL of 1× TFP Perm/Wash buffer. Permeabilization was achieved by resuspending the cells in 700 μL of Perm Buffer III (BD Biosciences) and incubating for 20 minutes at 4°C. Cells were then washed three times with 1 mL of 1× TFP Perm/Wash buffer and then resuspended in 50 μL of 1× TFP Perm/Wash buffer containing the following antibodies: FoxP3 APC (236A/E7, ThermoFisher Scientific 17-4777-42, 1 test volume/well), pSTAT5 PE (pTyr694, A17016B.Rec, BioLegend 936904, 1 test volume/well), and CD25 BV711 (M-A251, BioLegend 356138, 0.2 test volume/well). Cells were incubated at 4°C overnight. An additional 30uL of 1× TFP Perm/Wash buffer was added containing the following antibodies: CD3 FITC (UCHT1, BioLegend 300406, 0.2 test volume/well), CD4 BV421 (SK3 BioLegend 344632, 0.2 test volume/well), CD8 BV605 (SK1 BioLegend 344742, 0.2 test volume/well). Cells were incubated for 1 additional hour at 4°C and then washed twice with FACS buffer (a 1x DPBS (Thermo Fisher) solution with 2.5% FBS and 0.02% sodium azide). Data were collected on a Cytek Aurora flow cytometer and analyzed using FlowJo software. T_regs_ were gated as CD3^+^CD4^+^FOXP3^+^ cells, CD8^+^ T cells were gated as CD3^+^CD8^+^ cells, and T_conv_ cells were gated as CD3^+^CD4^+^FOXP3^-^ cells. pSTAT5 dose-response data were fitted to a nonlinear three-parameter response curve using GraphPad Prism v10. All experiments were conducted in triplicate unless otherwise noted.

## Appendix A. Supplementary material

Supplementary Figures S1–S18 and Tables 1-2.

Supplementary Videos 1-12.

## CRediT authorship contribution statement

CHW: conceptualization, methodology, formal analysis, investigation, writing – original draft, writing – review & editing, visualization; SOS: conceptualization, methodology, formal analysis, investigation, writing – original draft, writing – review & editing, visualization; DH: methodology, formal analysis, investigation, writing – review & editing; VSP: methodology, investigation, resources; CSF: methodology, resources– original draft; MCY: methodology, resources; AT: methodology, formal analysis, investigation, visualization; ET: investigation, resources; AM: methodology, resources, supervision; JBS: supervision, funding acquisition; GRB: supervision, funding acquisition; NGS: conceptualization, methodology, writing – original draft, writing – review & editing, supervision, funding acquisition.

## Data and Code Availability

NMR assignments have been deposited in the BMRB, under codes 52582 (S15), 52583 (S1), and 52591 (S1 L56A). All MD code relating to this paper is deposited at https://github.com/ssolieva/IL2-dynamics. Simulation trajectories are available upon request.

## Acknowledgments

This project was funded in part by the National Institute of Allergy and Infectious Diseases, National Institutes of Health, Department of Health and Human Services (NIH R01AI143997 to N.G.S.). Additional support was provided by the NexTGen team through the Cancer Grand Challenges partnership, funded by Cancer Research UK (CGCAT F-2021/100002), the National Cancer Institute (CA 278687-01), and the Mark Foundation for Cancer Research (to N.G.S.). The authors gratefully acknowledge a generous donation from the Asplundh Foundation that supported the purchase of SPR equipment used in this project. Further funding was provided by NSF MCB 2218156 and NIH NIGMS R35GM152085 (to G.R.B.), R35GM125034 (to N.G.S), which funded the cryoprobe of the 600 MHz NMR instrument at UPenn, as well as NIH NIBIB R01 EB029341, NSF CAREER 2143160, DoD W81XWH-21-1-0891 and HT94252310627, the Helmsley Charitable Trust, Kenneth Rainin Foundation, and Gabrielle’s Angel Foundation for Cancer Research (to J.B.S.). C.H.W. and S.O.S. were supported by T32GM132039. The authors would like to extend their heartfelt gratitude (in no particular order) to Dr. Possu Huang for sharing the plasmid constructs for S15 and S1, Drs. Thomas Malek and Milos Vujanac for sharing their pulse-chase IL-2 T cell stimulation protocol, Drs. Jinfa Ying and Ad Bax (NIH) for assistance with recording NMR data, Dr. Ryan Kubanoff and the Biological Chemistry Resource Center at UPenn for assistance with CD experiments, the Biomolecular NMR center at Johns Hopkins University for 800 MHz NMR time, the Flow Cytometry Core Laboratory at the Children’s Hospital of Philadelphia Research Institute for use of the flow cytometry instrumentation, Samuel Garfinkle for assistance with MD analysis, Yi Sun for assistance with surface plasmon resonance data collection, Sagar Gupta and Omar Ani for assistance with computational analysis of the CPMG data, Dr. Leland Mayne for advising the LC-MS/MS experiments, and Dr. Leena Mallik for initial crystallography screening of IL-2 variants. We thank the Human Immunology Core at the University of Pennsylvania for providing all T cells used in this study.

