## Supplementary Figures 1-18, Tables 1-2 for "Regulating IL-2 immune signaling function via a core allosteric structural network"

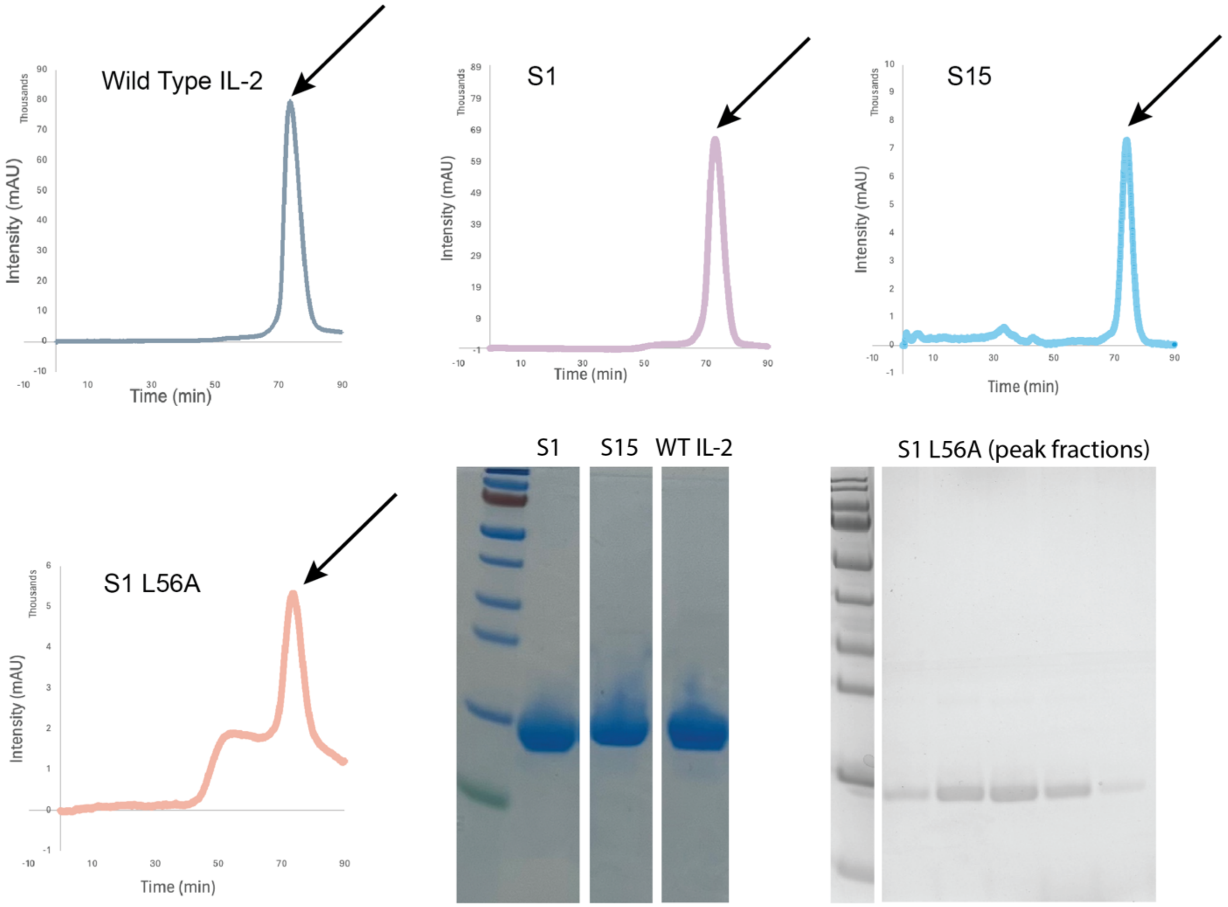


```
1          10                  20                  30
APTSSSTKKTQLQLEHLLLDLQMILNGINN
APTSSSTKKTQLQLEHLLLDLQMILNMLNN
APTSSSTKKTQLQLEHLLLDLQMILNMLNN

           40                  50                  60
YKNPKLTRMLTFKFYMPKKATELKHLQCLE
YDNPKLTRMLTFKFYMPKKATELKHLQCLE
YDNPKLTRLLTFKFYMPKKATSLKHLQCLE

           70                  80                  90
EELKPLEEVLNLAQSKNFHLRPRDLISNIN
EELKPLEEALNQAQSKNFHLDPRDLISNIN
EELKPLEEALNAAGDDPK--TIRDVVSNIN

           100                 110                 120
VIVLELKGSETTFMCEYADETATIVEFLNR
VIVLELKGSETTFMCEYADETATIIEFLNR
VFVLELKGSETTFMCEYADETATIIEFLNR

           130
WITFSQSIISTLT Wild Type
WITFSQSIISTLT S15
WITFSQSIISTLT S1
```

**Supplementary Figure 1: Purification of IL-2 variants.** Shown are size exclusion chromatography purification peaks and representative SDS-PAGE gels post-purification to demonstrate the high purity of the samples. A sequence alignment of the three proteins is shown at the bottom, with mutation sites colored red and highlighted sequence showing the alpha helical secondary structure, defined by crystal structures.

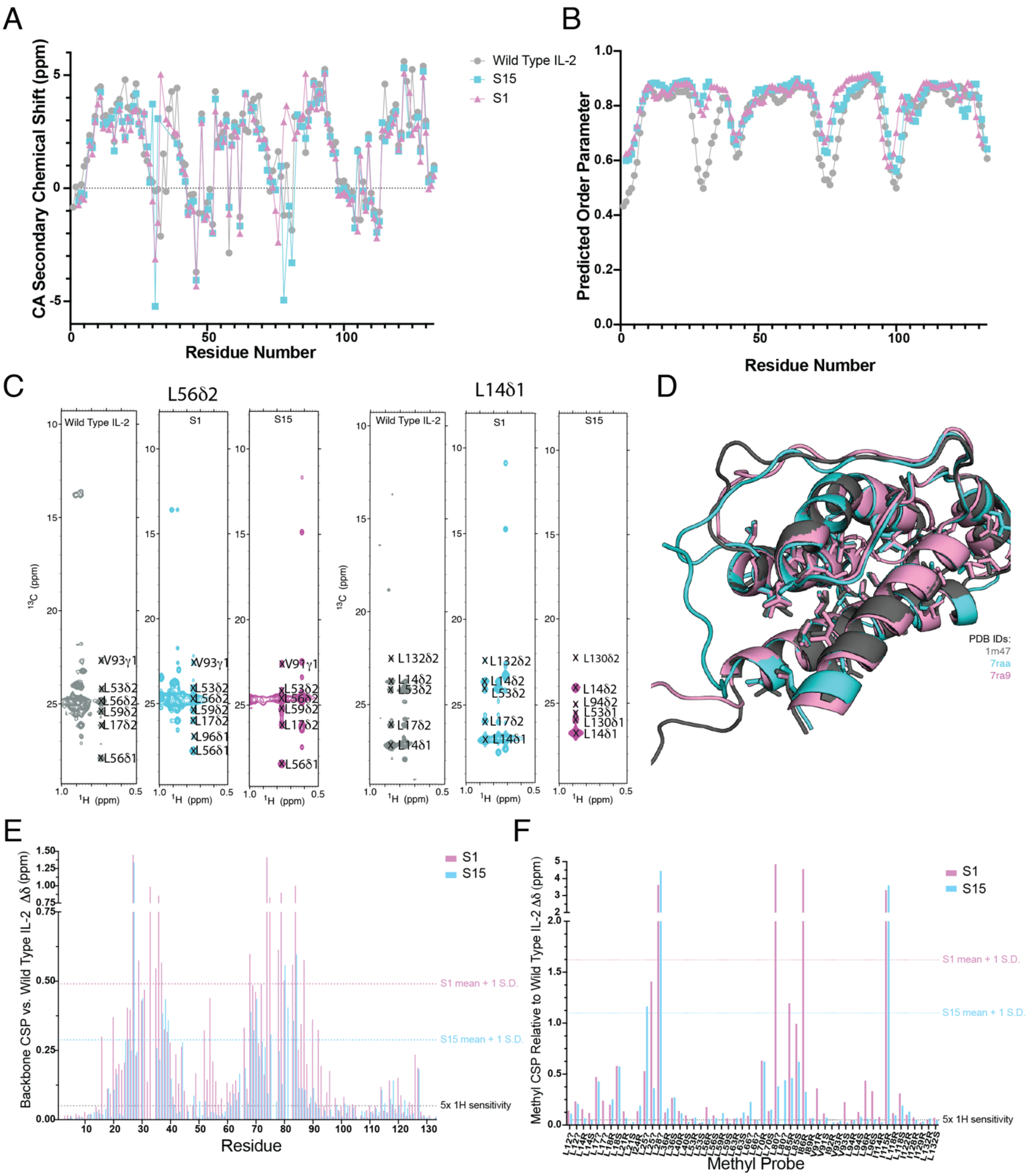


**Supplementary Figure 2: NMR assignment and calculated chemical shift perturbations.** A) CA secondary chemical shifts calculated from the TALOS-N software for wild type IL-2 (gray), S15 (cyan), and S1 (magenta). Generally, CA secondary chemical shifts are positive when alpha helical secondary structures are present. B) Predicted order parameters from TALOS-N, where very high values (>0.8) indicate rigid regions of the structure and lower values indicate local flexibility of the amide bond vector. C) 2D strips from Cm-CmHm NOESY data used to assign the isoleucine, leucine, and valine methyl peaks. D) Existing crystal structures of wild type IL-2 (gray), S15 (cyan), and S1 (magenta) overlayed with assigned methyls shown as sticks. Calculated chemical shift perturbations of E) backbone amide residues based on overlayed $^{15}$N TROSY HSQC spectra and F) ILE, Leu, and Val methyl residues based on methyl $^{13}$C SOFAST HMQC spectra, all collected at 800 MHz magnetic field. Both S1 (magenta) and S15 (cyan) were calculated relative to the wild type IL-2 spectra. All residue numbering is based on alignment to the wild type sequence.

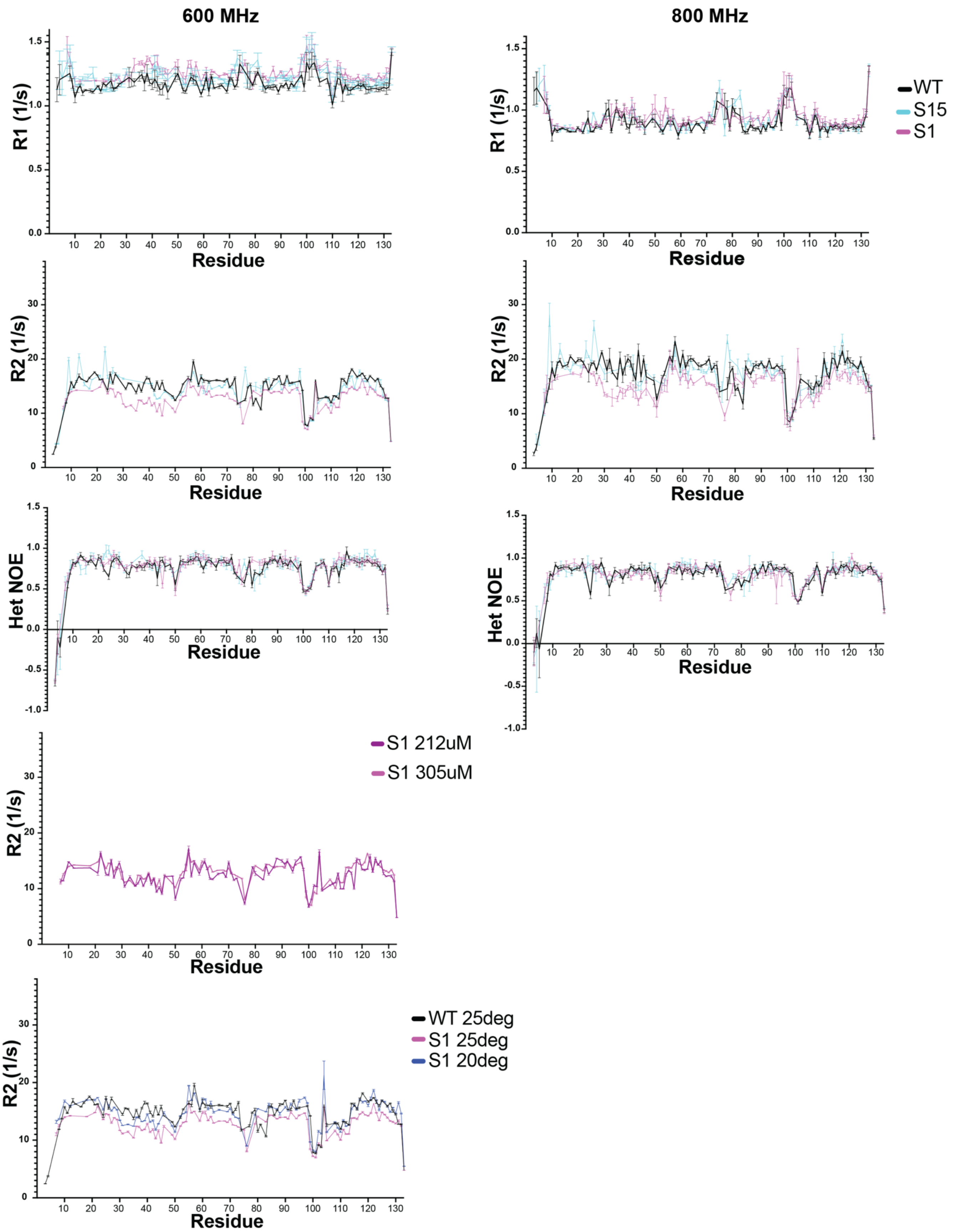


**Supplementary Figure 3: Spin relaxation datasets derived from $R_1$, $R_2$, and heteronuclear NOE experiments**. Shown are data collected at 600 MHz and 800 MHz magnetic fields for wild type IL-2 (black), S1 (magenta), and S15 (cyan). Given the clear baseline shift in the $R_2$ data for S1, $R_2$ experiments for S1 at different concentrations and temperatures are also shown in the bottom left panels. All residue numbering is based on alignment to the wild type sequence.

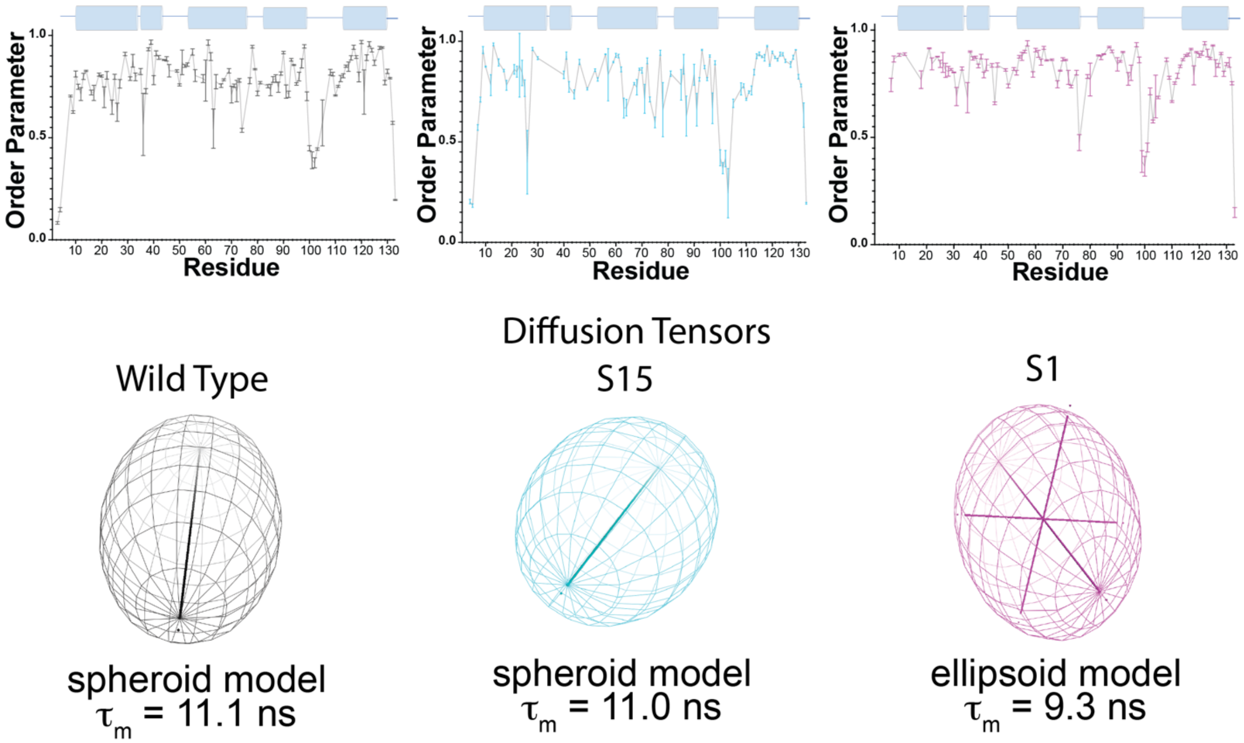


**Supplementary Figure 4: Order parameters calculated from NMR relaxation experiments.** Experimental $R_1$, $R_2$, and heteronuclear NOE datasets collected at both 600 MHz and 800 MHz magnetic fields were used to calculate order parameters using the Relax model-free software, shown in the top panels for wild type IL-2 (left), S15 (middle) and S1 (right) with secondary structure plotted above for reference. Fitted diffusion tensors and correlation times ($\tau_m$) are shown below. All residue numbering is based on alignment to the wild type sequence.

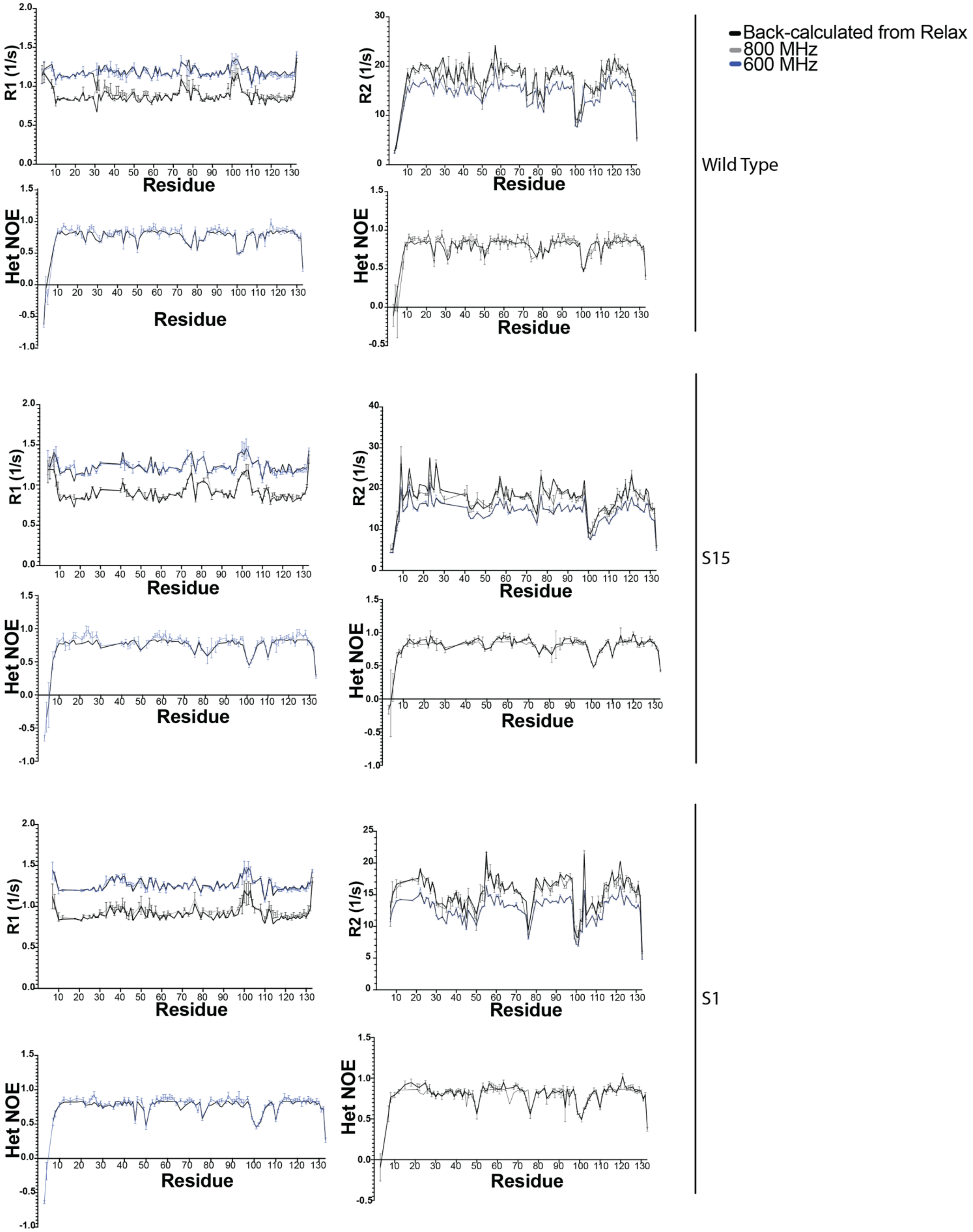


**Supplementary Figure 5: Back-calculated spin relaxation data based on the rotational diffusion models.** In the Relax model-free program, back-calculated $R_1$, $R_2$, and heteronuclear NOE values can be generated from the fitted models to verify how well the experimental data can accurately be described by the models. Back-calculated data are plotted as a black line, with raw data shown in blue (600 MHz) or gray (800 MHz). All residue numbering is based on alignment to the wild type sequence.

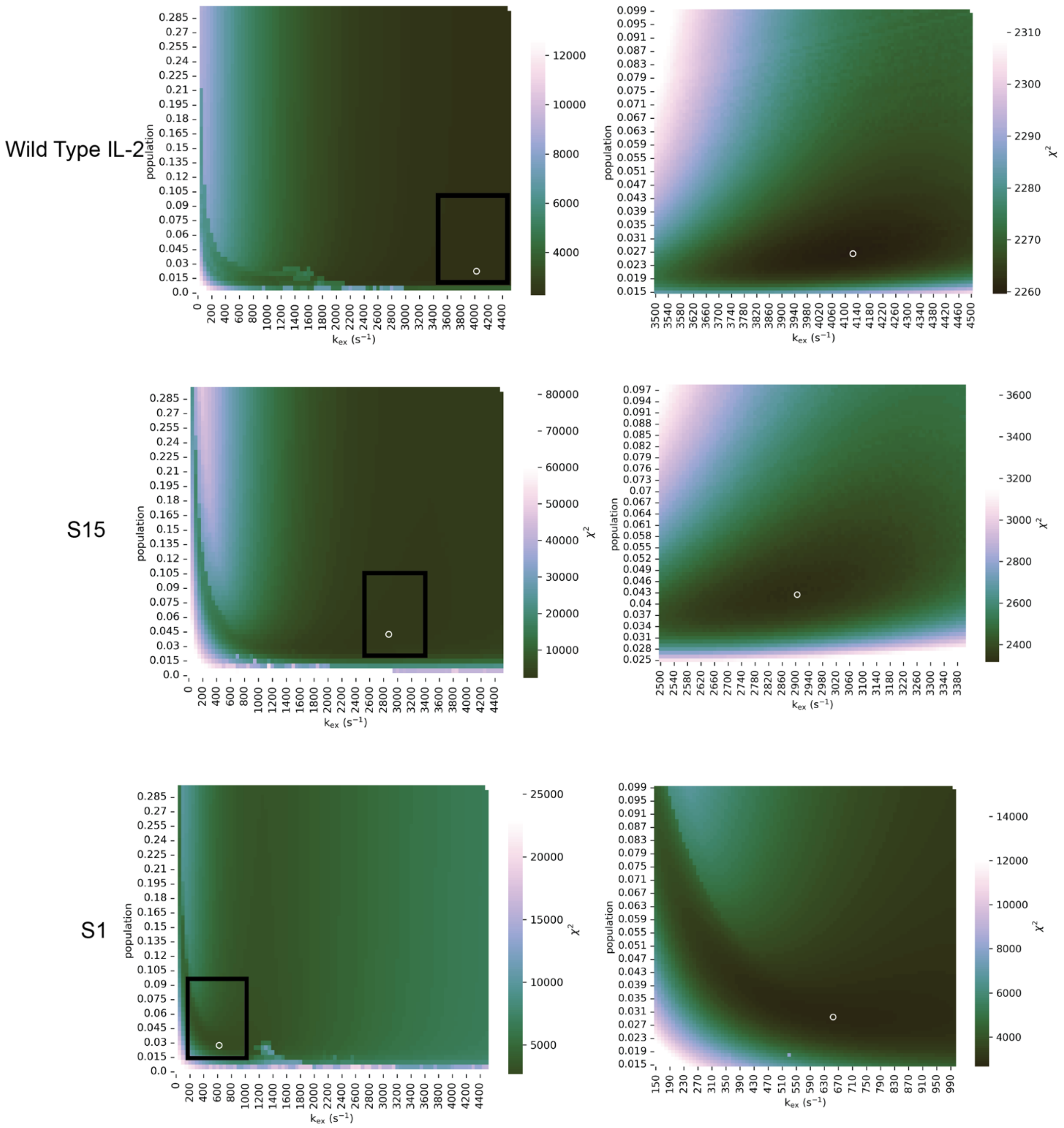


**Supplementary Figure 6*:* $\chi^2$ surface plots.** Surface plots were generated from fits of CPMG relaxation dispersion data recorded at 600 and 800 MHz magnetic fields on $^{13}C$ methyl labeled IL-2 variants (top, wild type IL-2; middle, S15; bottom, S1). The panels on the right represent a refined search near the global minimum, represented by the black squares on the left panels. To generate these plots, CPMG relaxation dispersion data was iteratively fitted to the two-site exchange model using the CATIA software, with kex and population (pb) held constant during each run, and the $\chi^2$ value representing the goodness of fit was extracted. The |Δw| values were not fixed during the search for the minimum $\chi^2$ value. In each panel, the white circle indicates the position of (pb, kex) that corresponds to the global minimum of $\chi^2$.

Wild Type IL-2 residues included in final global fit:

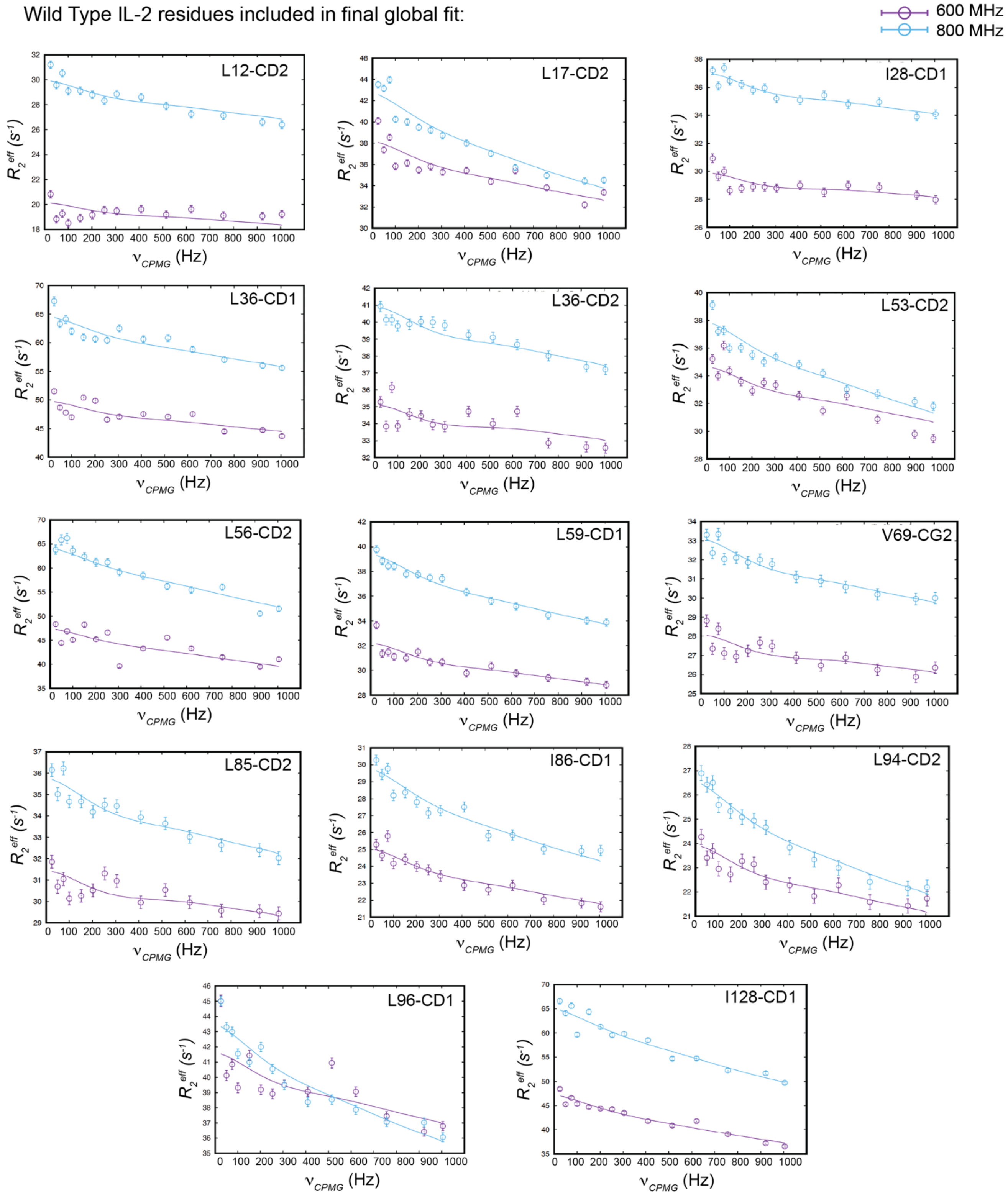


**Supplementary Figure 7: Dispersion curves for all exchanging residues in wild type IL-2.** CPMG relaxation dispersion data were collected at 600 MHz (purple) and 800 MHz (blue) magnetic fields at 5°C. Experimental relaxation dispersion profiles (circles) for residues exhibiting μs-ms timescale dynamics as measured by $^{13}$C single quantum CPMG relaxation dispersion experiments. Solid lines represent the best fit to a global two-site exchange model. These residues were used for the global analysis using the CATIA program.

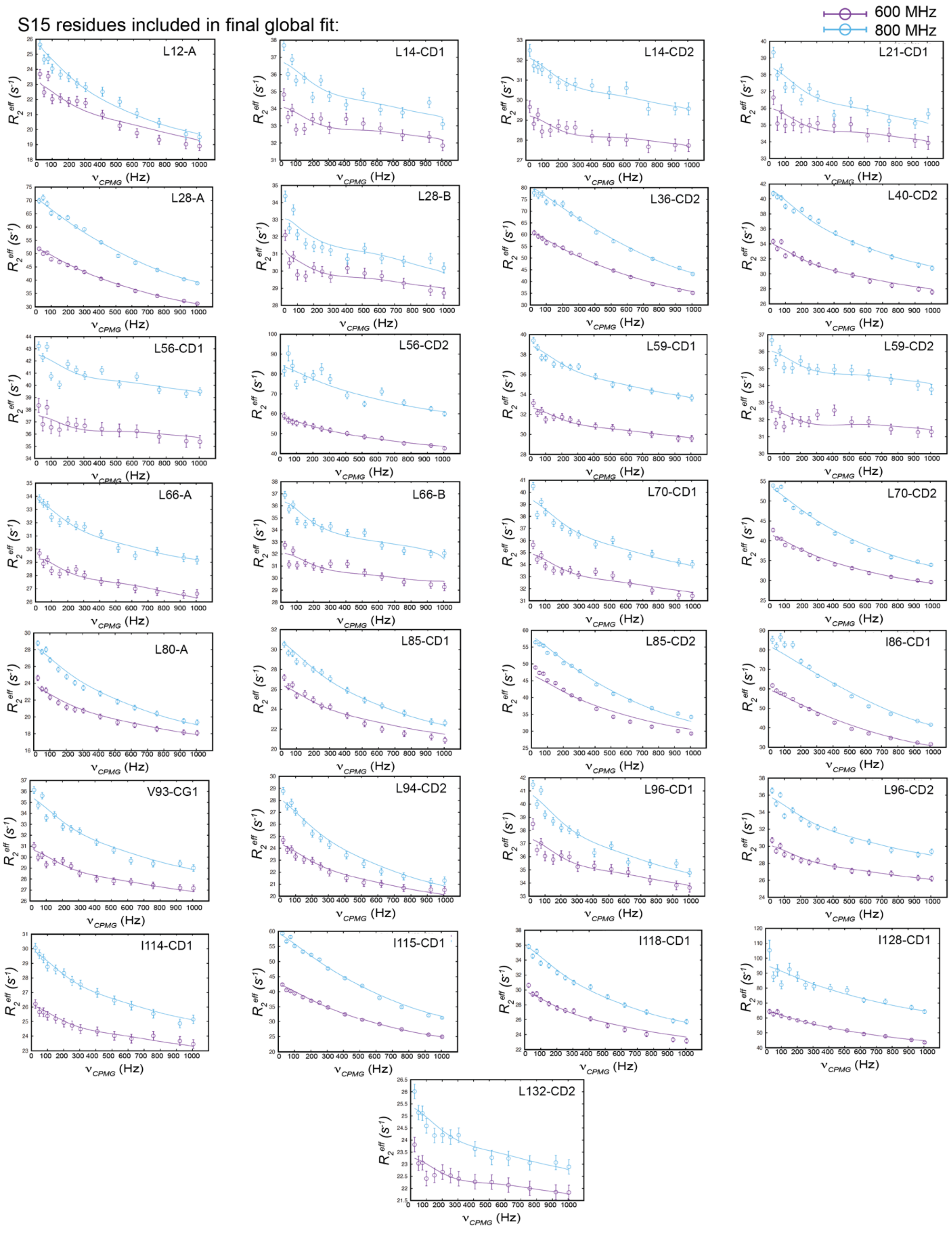


**Supplementary Figure 8: Dispersion curves for all exchanging residues in the S15 superkine.** CPMG relaxation dispersion data were collected at 600 MHz (purple) and 800 MHz (blue) magnetic fields at 5°C. Experimental relaxation dispersion profiles (circles) for residues exhibiting μs-ms timescale dynamics as measured by $^{13}$C single quantum CPMG relaxation dispersion experiments. Solid lines represent the best fit to a global two-site exchange model. These residues were used for the global analysis using the CATIA program.

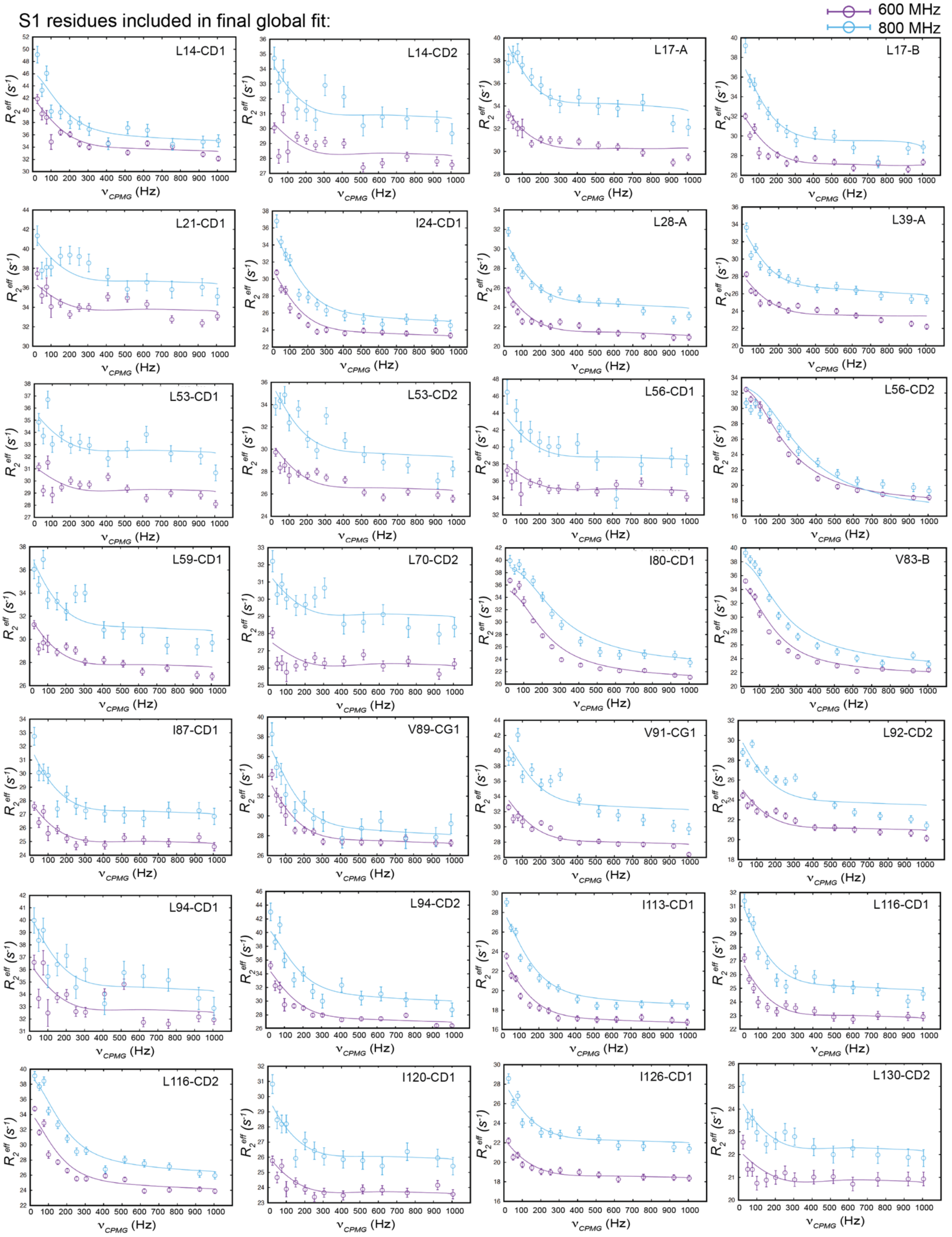


**Supplementary Figure 9: Dispersion curves for all exchanging residues in the S1 superkine.** CPMG relaxation dispersion data were collected at 600 MHz (purple) and 800 MHz (blue) magnetic fields at 5°C. Experimental relaxation dispersion profiles (circles) for residues exhibiting µs-ms timescale dynamics as measured by 13C single quantum CPMG relaxation dispersion experiments. Solid lines represent the best fit to a global two-site exchange model. These residues were used for the global analysis using the CATIA program.

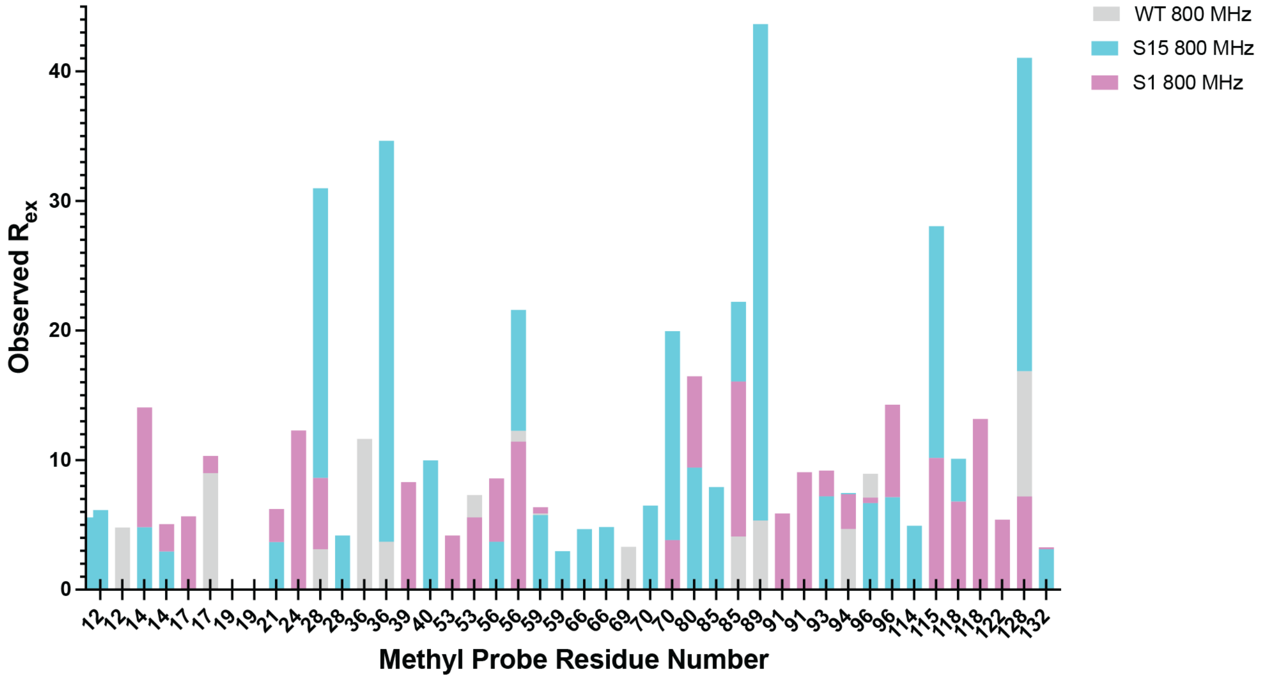


Mean: 7.22
Median: 5.63
Standard Deviation: 4.22

Mean: 12.63
Median: 7.16
Standard Deviation: 12.10

Mean: 8.64
Median: 7.83
Standard Deviation: 3.68

**Supplementary Figure 10: Plots of the $R_{ex}$ contributions of the methyl groups.** $R_{ex}$ contributions were calculated from the differences between $R_2^{eff}$ (25 Hz) and $R_2^{eff}$ (1005 Hz). The methyl groups with significant $R_{ex}$ contributions (> 3 Hz) are colored gray for wild type IL-2, cyan for S15, and magenta for S1. Shown below the chart are the mean, median, and standard deviation metrics for each of the three proteins. All residue numbering is based on alignment to the wild type sequence.

**Supplementary Table 1: Data summary from the CPMG relaxation dispersion experiments and global two-site exchange fits.** The upper table summarizes the global fitted parameters for each protein. Below are the residues included in each fit along with their fitted |Δω| values and errors.

| | # Methyls Fitted | Fitted $p_B$ | Fitted $k_{ex}$ ($s^{-1}$) |
|---|---|---|---|
| Wild Type IL-2 | 14 | 0.027 ± 0.005 | 4088 ± 190 |
| S15 | 29 | 0.043 ± 0.001 | 2878 ± 44 |
| S1 | 28 | 0.030 ± 0.001 | 651 ± 16 |

Wild Type IL-2

| Residue | \|Δω\| | Error |
|---|---|---|
| L12D2 | 0.631 | 0.058 |
| L17D2 | 1.159 | 0.109 |
| I28D1 | 0.604 | 0.056 |
| L36D1 | 1.120 | 0.105 |
| L36D2 | 0.661 | 0.060 |
| L53D2 | 0.986 | 0.090 |
| L56D2 | 1.389 | 0.137 |
| L59D1 | 0.884 | 0.079 |
| V69G2 | 0.658 | 0.060 |
| L85D2 | 0.676 | 0.061 |
| I86D1 | 0.870 | 0.078 |
| L94D2 | 0.795 | 0.071 |
| L96D1 | 1.048 | 0.07 |
| I128D1 | 1.616 | 0.169 |

S15

| Residue | \|Δω\| | Error |
|---|---|---|
| L12A | 0.567 | 0.013 |
| L14D1 | 0.402 | 0.015 |
| L14D2 | 0.358 | 0.016 |
| L21D1 | 0.407 | 0.016 |
| L28A | 1.537 | 0.032 |
| L28B | 0.379 | 0.016 |
| L36D2 | 1.754 | 0.040 |
| L40D2 | 0.763 | 0.014 |
| L56D1 | 0.391 | 0.017 |
| L56D2 | 1.237 | 0.040 |
| L59D1 | 0.523 | 0.013 |
| L59D2 | 0.302 | 0.019 |
| L66A | 0.490 | 0.014 |
| L66B | 0.475 | 0.014 |
| L70D1 | 0.539 | 0.013 |
| L70D2 | 1.124 | 0.020 |
| L80A | 0.715 | 0.014 |
| L85D1 | 0.683 | 0.013 |
| L85D2 | 1.315 | 0.025 |
| I86D1 | 1.928 | 0.045 |
| V93G1 | 0.597 | 0.013 |
| L94D2 | 0.630 | 0.013 |
| L96D1 | 0.566 | 0.013 |
| L96D2 | 0.610 | 0.013 |
| I114D1 | 0.502 | 0.013 |
| I115D1 | 1.394 | 0.027 |
| L118D1 | 0.764 | 0.014 |
| I128D1 | 1.494 | 0.041 |
| L132D2 | 0.361 | 0.016 |

S1

| Residue | \|Δω\| | Error |
|---|---|---|
| L14D1 | 0.590 | 0.024 |
| L14D2 | 0.256 | 0.017 |
| L17A | 0.324 | 0.016 |
| L17B | 0.428 | 0.013 |
| L21D1 | 0.291 | 0.022 |
| I24D1 | 0.544 | 0.014 |
| L28A | 0.369 | 0.010 |
| L39A | 0.391 | 0.011 |
| L53D1 | 0.230 | 0.017 |
| L53D2 | 0.362 | 0.015 |
| L56D1 | 0.311 | 0.027 |
| L56D2 | 1.202 | 0.027 |
| L59D1 | 0.362 | 0.015 |
| L70D2 | 0.195 | 0.017 |
| I80D1 | 1.147 | 0.027 |
| V83B | 0.898 | 0.019 |
| I87D1 | 0.294 | 0.014 |
| V89G1 | 0.474 | 0.019 |
| V91G1 | 0.477 | 0.015 |
| L92D2 | 0.380 | 0.010 |
| L94D1 | 0.343 | 0.021 |
| L94D2 | 0.561 | 0.019 |
| I113D1 | 0.499 | 0.011 |
| L116D1 | 0.363 | 0.011 |
| L116D2 | 0.690 | 0.015 |
| I120D1 | 0.258 | 0.014 |
| I126D1 | 0.340 | 0.011 |
| L130D2 | 0.189 | 0.013 |

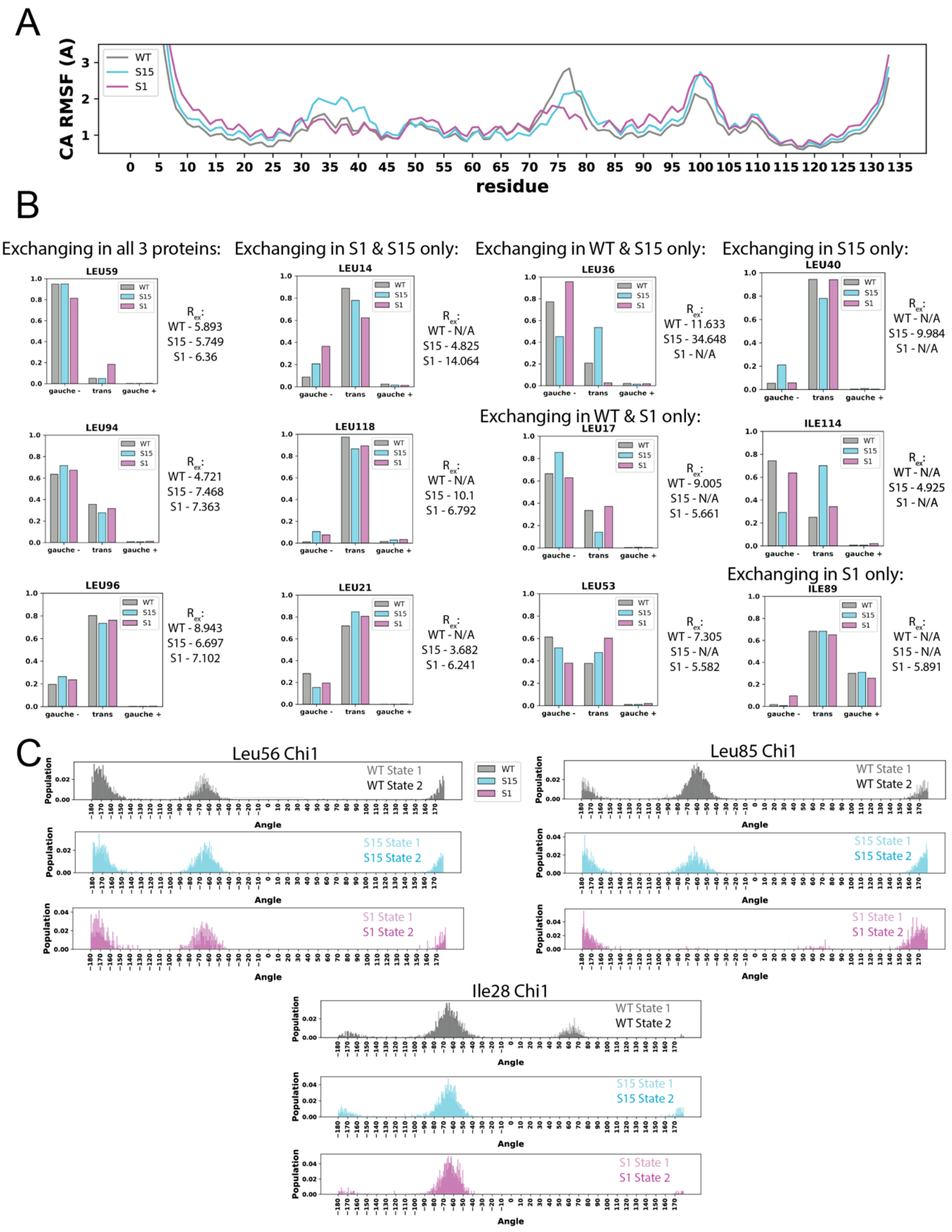


**Supplementary Figure 11: Qualitative comparison between intermediate-timescale dynamics by CPMG NMR and structural plasticity by MD.** A) Average alpha carbon root-mean-square fluctuation (RMSF) shows similar backbone fluctuations between the three proteins, with S15 having more fluctuations in the mini helix region. B) Residues with significantly varied $\chi_2$ angle distributions across the three proteins in the simulations are shown as bar graphs. We note that the residues with $\chi_2$ angle differences in the simulations are the same residues undergoing conformational exchange by NMR, and in most cases the population distributions roughly follow the trends of estimated $R_{ex}$ variations across proteins (shown here as measured at 800 MHz $^1$H magnetic field). C) When clustered into "major" and "minor" states using time-lagged independent component analysis (tICA), we observe multiple examples of residues with high $R_{ex}$ in NMR showing detectable shifts in Chi1 angle distribution across states in MD frames. All residue numbering is based on alignment to the wild type sequence.

**Table 2: Input residues and atoms for tICA pairwise distance analyses.** Residue numbering is based on alignment to the wild type sequence.

| Residue ID, Atom |
|---|
| LEU12, CD1 |
| LEU14, CD1 |
| LEU17, CD1 |
| LEU18, CD1 |
| LEU19, CD1 |
| LEU21, CD1 |
| ILE24, CD1 |
| LEU25, CD1 |
| LEU36, CD1 |
| LEU40, CD1 |
| LEU53, CD1 |
| LEU59, CD1 |
| LEU63, CD1 |
| LEU66, CD1 |
| LEU70, CD1 |
| ILE89, CD1 |
| VAL91, CG1 |
| VAL93, CG1 |
| LEU94, CD1 |
| LEU96, CD1 |
| ILE114, CD1 |
| LEU118, CD1 |
| ILE122, CD1 |
| ILE128, CD1 |
| ILE129, CD1 |
| LEU132, CD1 |

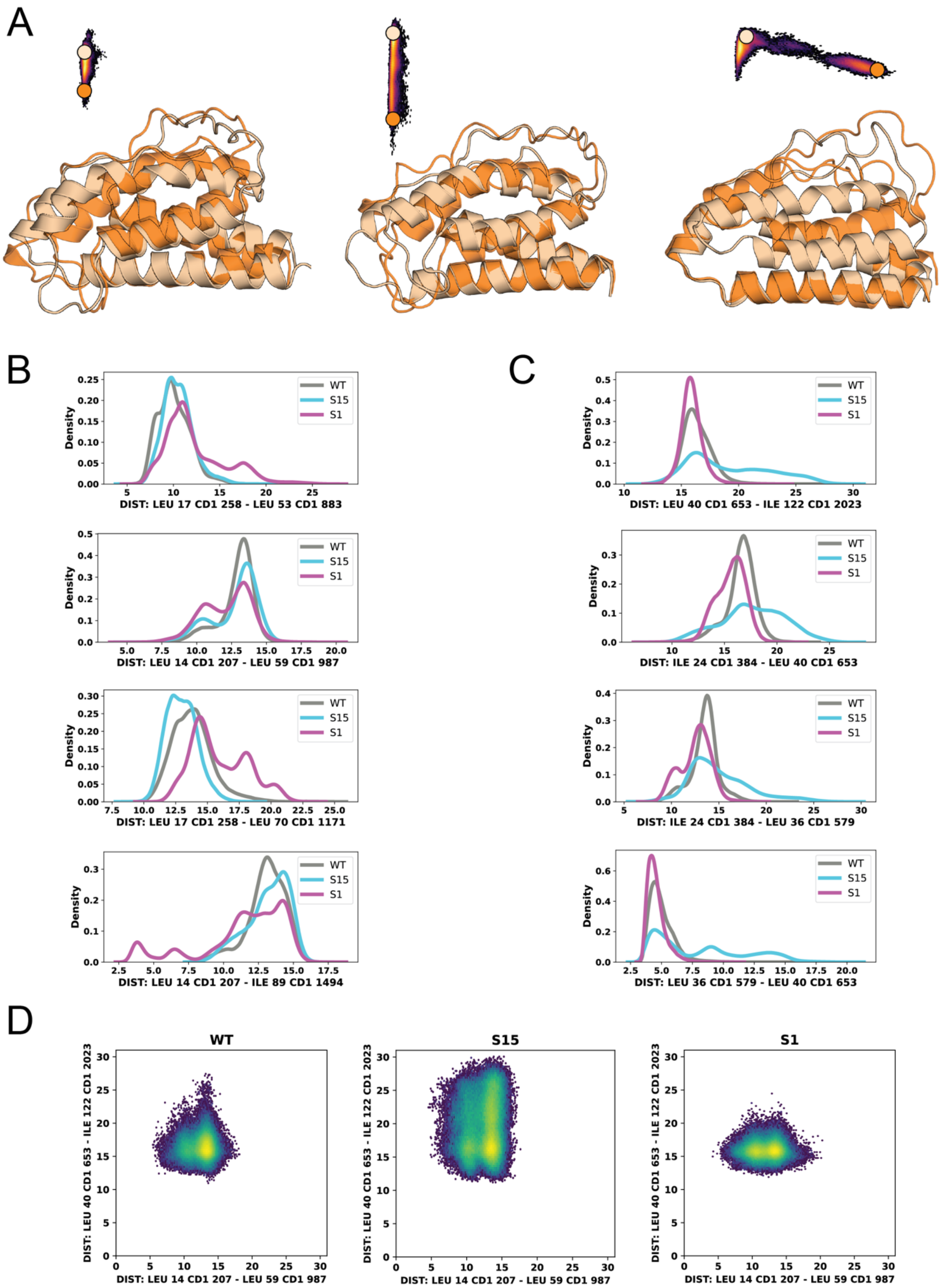


**Supplementary Figure 12: Comparing WT and superkines using time-lagged independent component analysis.** A) Overlays of representative frames from the "major" (tan) and "minor" (orange) conformations, also represented as colored circles on tICA projections for reference, for wild type IL-2 (left), S15 (center), and S1 (right). B) Distributions of the top 4 pairwise distances contributing to independent component 1 (IC1). C) Distributions of the top 4 pairwise distances contributing to independent component 2 (IC2). D) 2D density plots showing frame distributions across two key pairwise distances, L14-L59 and L40-I122. These report of movement and bending of the B-helix, a distinguishing motion across the three proteins. All residue numbering is based on alignment to the wild type sequence.

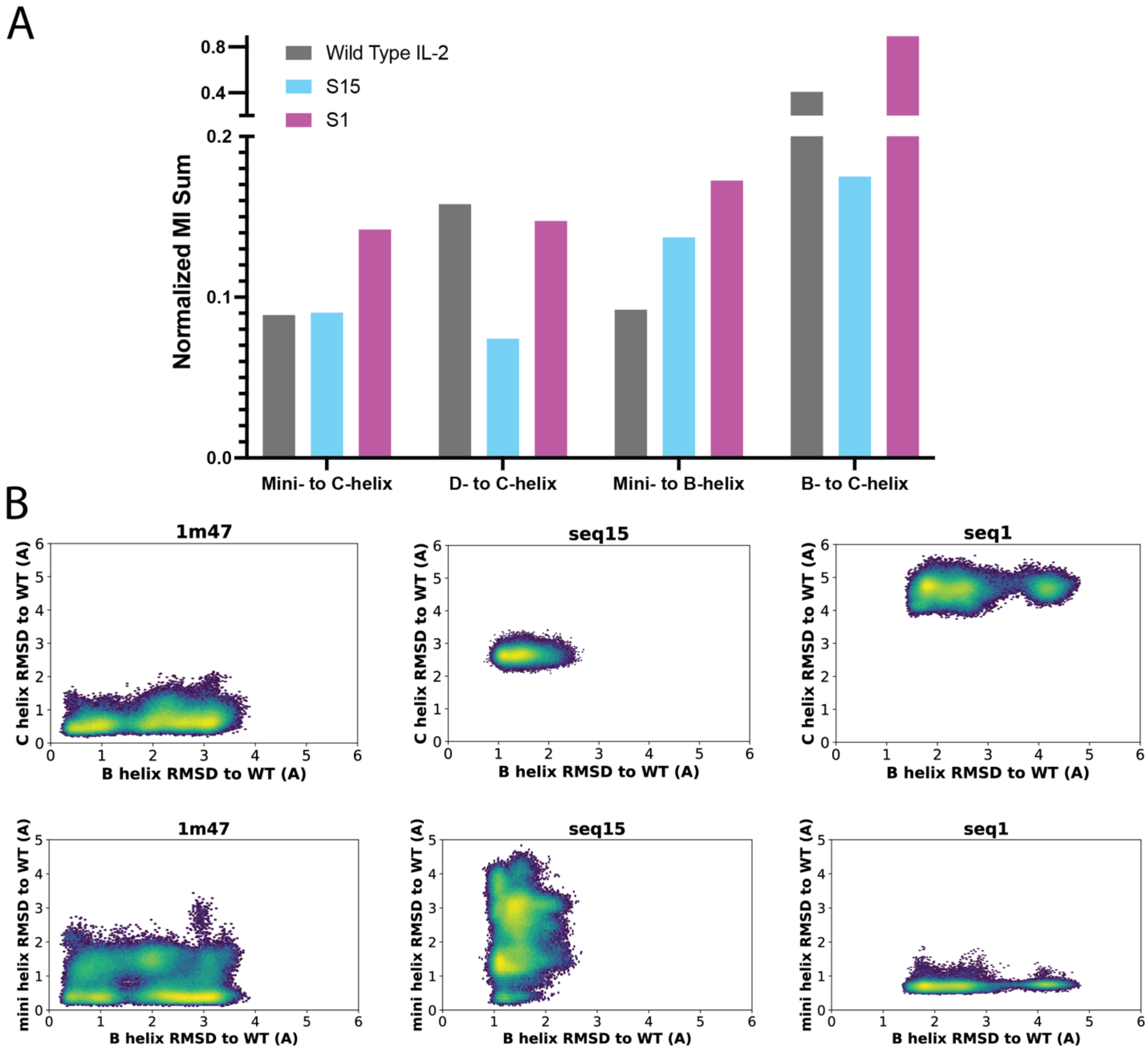


**Supplementary Figure 13: CARDS mutual information scores reveal allosteric correlations between key regions of IL-2 variants.** A) Normalized MI sums across key structural regions on IL-2 and superkines. MI scores were normalized based on the number of residues in a given region, as this varies across the three proteins. B) 2D RMSD plots of wild type IL-2 (1m47), S15 (seq15), and S1 (seq1) calculated relative to the starting frame in the wild type IL-2 simulation.

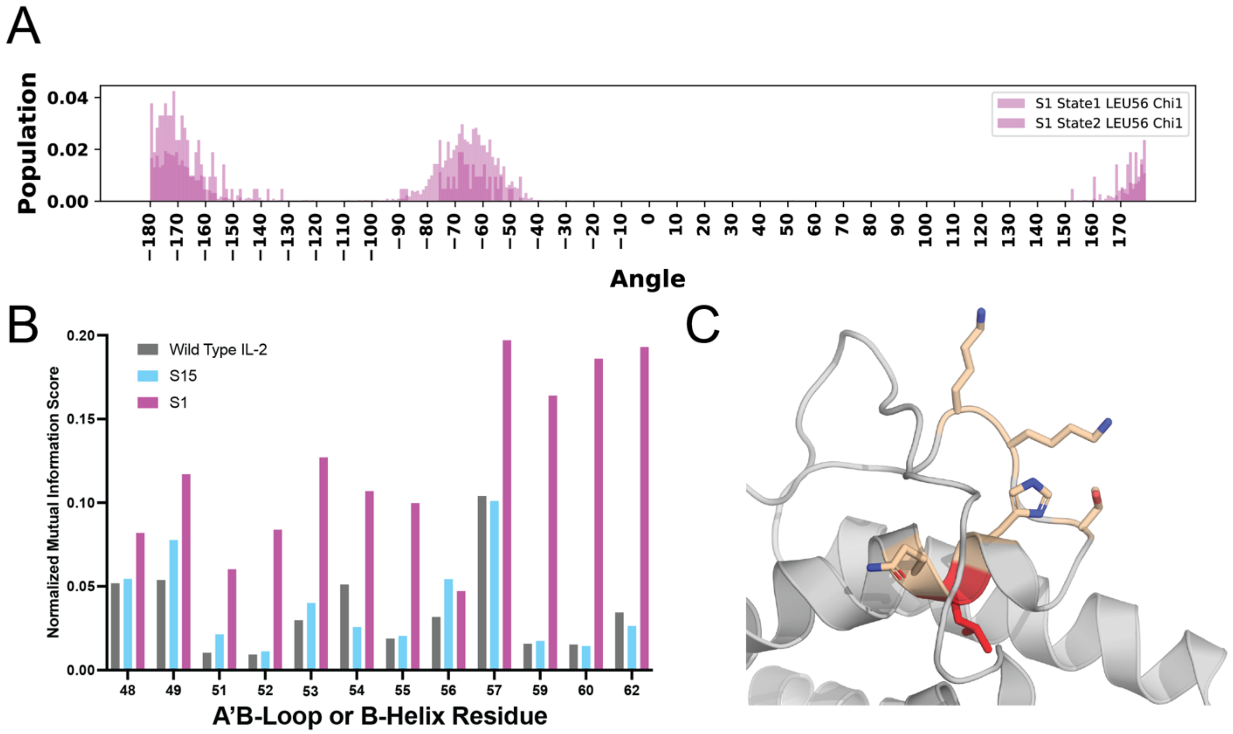


**Supplementary Figure 24: L56A mutation is predicted to disrupt the S1 allosteric network by MD simulations.** A) When clustered into “major” and “minor” states using time-lagged independent component analysis (tICA), we observe a strong switch in Leu56 in Chi1 angle distribution across “major” and “minor” states in S1 MD frames, in line with our with high $R_{ex}$ in NMR. B) The normalized sum of the mutual information values between a given residue and all other residues in the protein for wild type IL-2, S15, and S1, focused only on residues from the key cluster in the S1 allosteric network (resid. 48-62). C) Residues with the highest normalized sum of mutual information values in S1 are all near each other in the structure. Leu56 is central to these residues and is on the B helix.

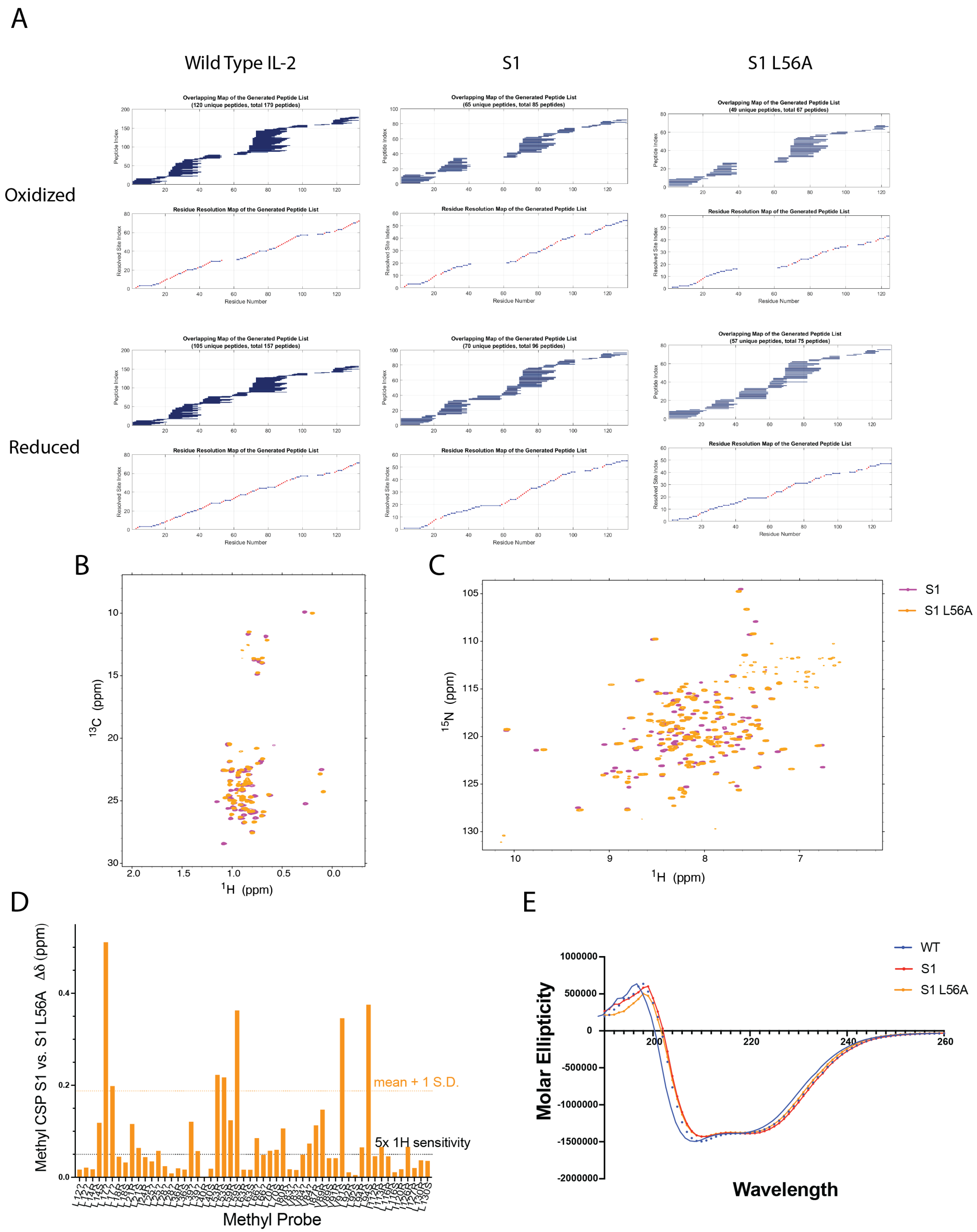


**Supplementary Figure 15. Experimental characterization of S1 L56A mutant in solution. A**) LC-MS/MS data demonstrating that wild type IL-2, S1, and S1 L56A proteins all have missing peptide reads at the site of the disulfide bridge when proteins are digested from the oxidized states. As the software used for these experiments cannot handle disulfide bond detection, reads could only be recovered at these sites when proteins were reduced with 1mM DTT prior to pepsin digestion, suggesting that all three proteins form the disulfide bridge in the oxidized state as expected. B) Overlayed $^{13}$C-$^{1}$H SOFAST methyl HMQC spectra and C) $^{15}$N-$^{1}$H TROSY HSQC spectra collected at 800 MHz magnetic field at 25°C for S1 (magenta) and S1 L56A (orange) proteins. D) Calculated methyl chemical shift perturbations (CSPs) for S1 L56A relative to S1. Dotted lines represent standard metrics for significance: mean + standard deviation (orange dotted line) and 5x $^{1}$H sensitivity of NMR instrument (black dotted line). E) Circular dichroism wavelength scans of 7μM samples of wild type IL-2 (blue), S1 (red), and S1 L56A (orange) from 260-190 nm at 25°C, in concentration-corrected molar ellipticity units.

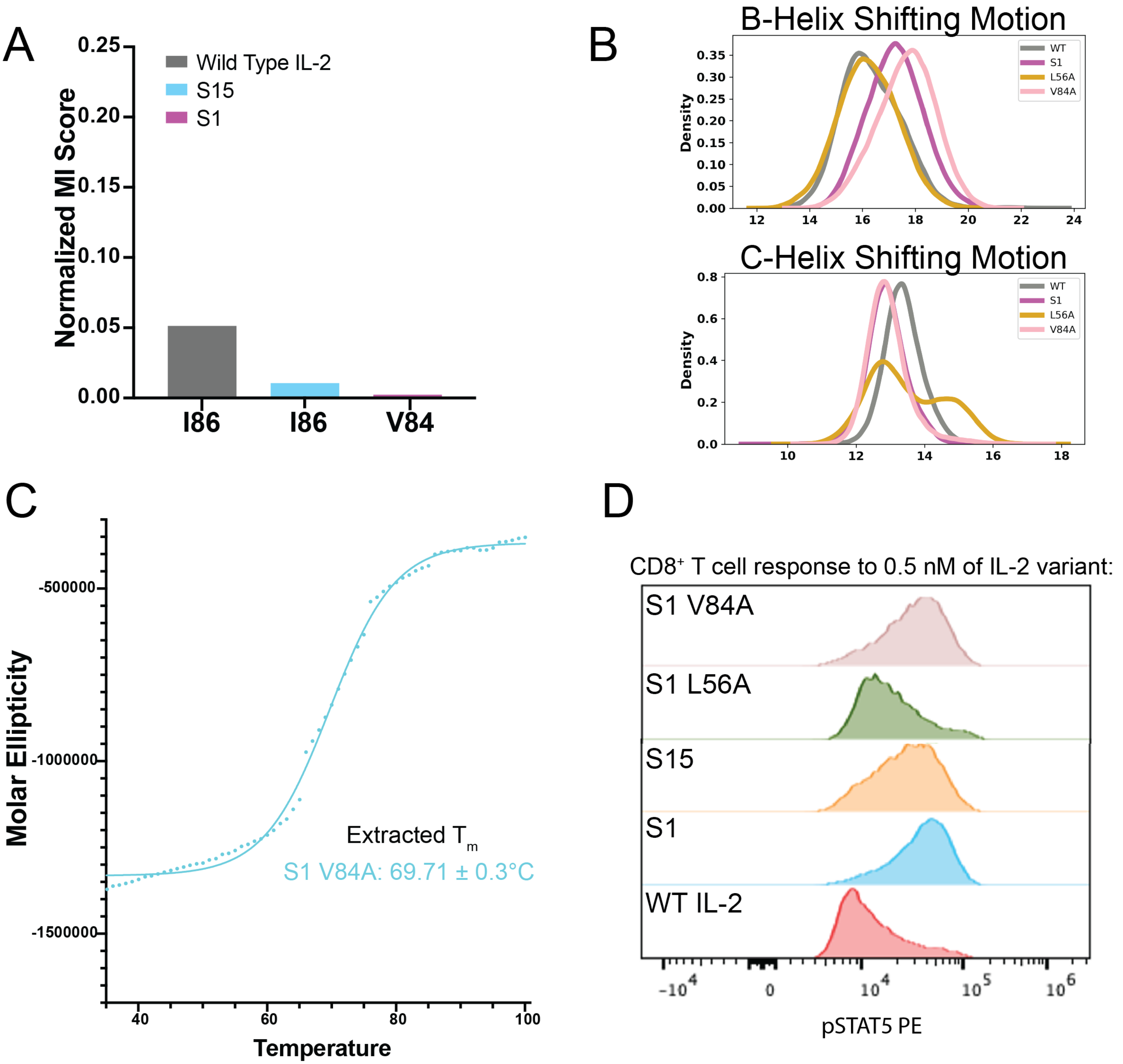


**Supplementary Figure 16: The S1 V84A mutant demonstrates that reduced thermal stability alone does not necessarily disrupt dynamics or allostery.** MD simulations and experiments of S1 V84A show that it does not disrupt the allosteric network in S1. A) Residue 84 has a very low mutual information score based on CARDS output for S1. B) Average distance distributions between residues of interest show that the dynamics of S1 V84A are extremely similar to S1 in MD simulations. For the B-helix shifting distribution, the following pairwise distances were averaged: L12-L70, L18-L66, L14-L59, L21-L70. For the C-helix shifting distribution, the following pairwise distances were averaged: L18-L96, L18-V91, L17-V91, L14-V91. C) Circular dichroism thermal melt curve for S1 V84A collected at 222 nm in 1°C intervals from 25°C to 98°C with extracted $T_m$ values fitted from a nonlinear Boltzmann sigmoidal curve fit in GraphPad Prism. D) STAT5 phosphorylation (pSTAT5) response via phospho-flow cytometry to a 0.5 nM dose of wild type IL-2, S15, S1, S1 L56A, or S1 V84A treatment in human donor $CD8^+$ T cells, demonstrating whether a given variant is more 'wild type-like' or 'superkine-like' in signal response. Pre-isolated total T cells from human donors were incubated with IL-2 variant for 20 min at 37°C prior to fixation, permeabilization, and staining.

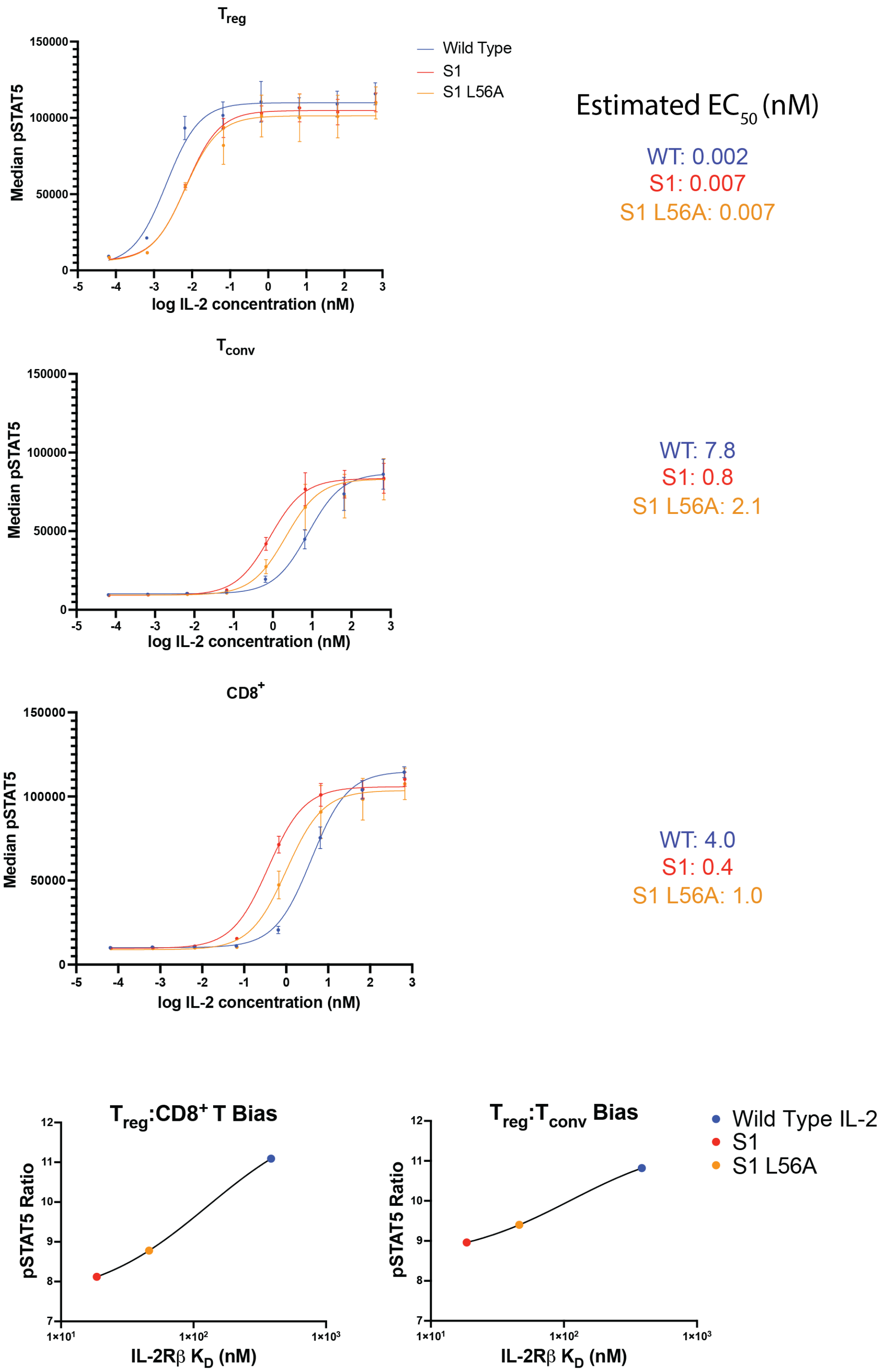


**Supplementary Figure 17: T cell signaling responses to IL-2 variants from a second human donor sample.** Shown are STAT5 phosphorylation responses to IL-2 in FoxP3⁺ $T_{reg}$ (top), CD4⁺ $T_{conv}$ (middle), and CD8⁺ $T_{eff}$ (bottom) cells from fresh human donor total T cells. T cells were incubated with IL-2 variant for 20 min at 37 °C. Dose-response curves were fitted to nonlinear three-parameter agonist vs. response curves in GraphPad Prism to extract $EC_{50}$ values shown on the right. Bottom: $T_{reg}$:CD8⁺ T and $T_{reg}$:$T_{conv}$ pSTAT5 ratios for IL-2 variants at 50 pM concentration are plotted against IL2Rβ $K_D$ (nM).

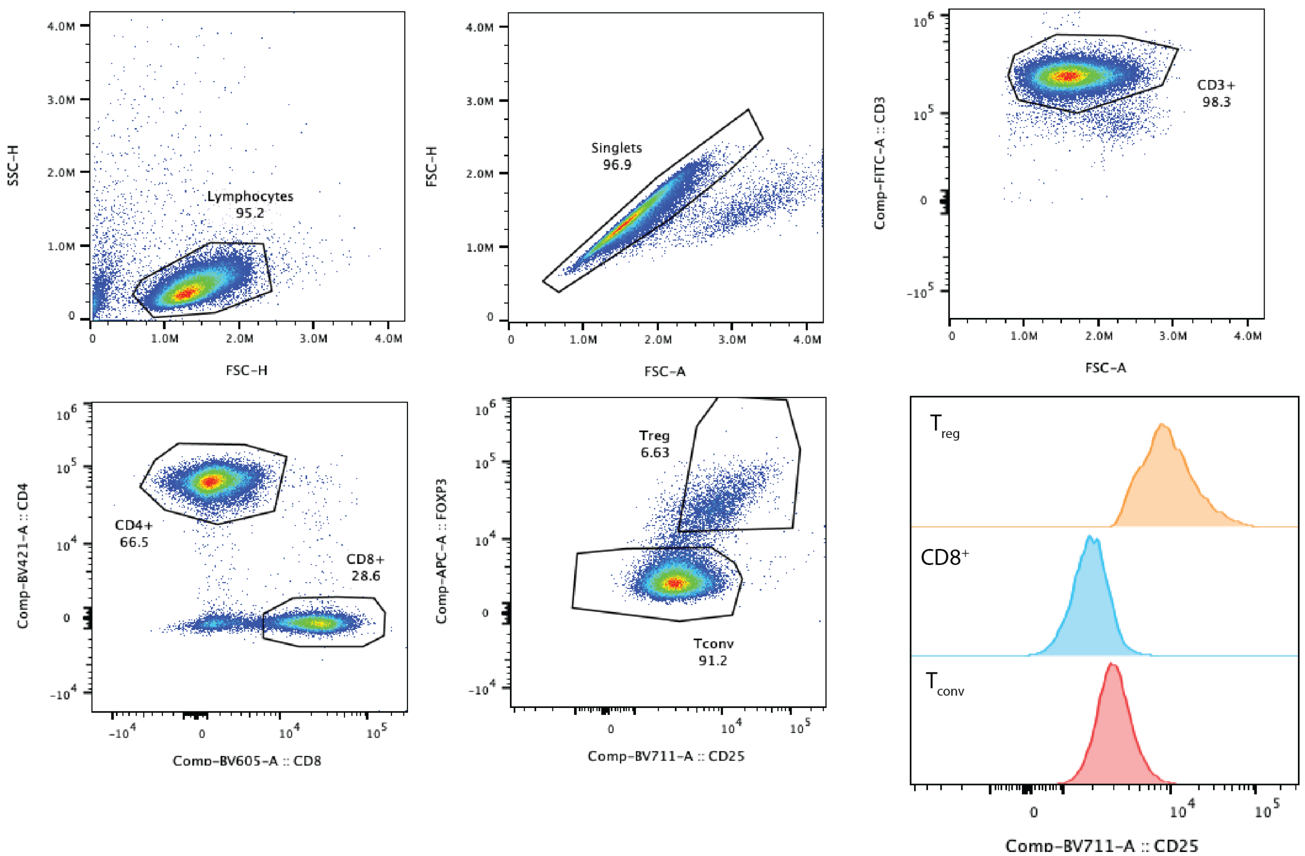


**Supplementary Figure 18: Gating strategy for phospho-flow cytometry-based pSTAT5 assays**. $T_{conv}$ cells were gated as Lymphocytes > Singlets > $CD3^+$ > $CD4^+$ > $FoxP3^-$. $T_{reg}$ cells were gated as Lymphocytes > Singlets > $CD3^+$ > $CD4^+$ > $FoxP3^+$. CD8+ T cells were gated as Lymphocytes > Singlets > $CD3^+$ > $CD8^+$. The bottom right corner depicts expression of CD25 in each T cell type, which is significantly higher in $T_{reg}$ cells than in either $T_{conv}$ or $CD8^+$ T cells, as expected.
